# Parsing Brain Network Specialization: A Replication and Expansion of Wang et al. (2014)

**DOI:** 10.1101/2024.02.13.580153

**Authors:** Madeline Peterson, Dorothea L. Floris, Jared A. Nielsen

## Abstract

One organizing principle of the human brain is hemispheric specialization, or the dominance of a specific function or cognitive process in one hemisphere or the other. Previously, Wang et al. (2014) identified networks putatively associated with language and attention as being specialized to the left and right hemispheres, respectively; and a dual-specialization of the executive control network. However, it remains unknown which networks are specialized when specialization is examined within individuals using a higher resolution parcellation, as well as which connections are contributing the most to a given network’s specialization. In the present study, we estimated network specialization across three datasets using the autonomy index and a novel method of deconstructing network specialization. After examining the reliability of these methods as implemented on an individual level, we addressed two hypotheses. First, we hypothesized that the most specialized networks would include those associated with language, visuospatial attention, and executive control. Second, we hypothesized that within-network contributions to specialization would follow a within-between network gradient or a specialization gradient. We found that the majority of networks exhibited greater within-hemisphere connectivity than between-hemisphere connectivity. Among the most specialized networks were networks associated with language, attention, and executive control. Additionally, we found that the greatest network contributions were within-network, followed by those from specialized networks.

**Significance Statement:** Hemispheric specialization is a characteristic of brain organization that describes when a function draws on one hemisphere of the brain more than the other. We sought to identify the most specialized brain networks within individuals, as well as which connections contribute the most to a given network’s specialization. Among the most specialized networks were those associated with language, attention, and executive control. Unexpectedly, we also identified networks associated with emotion/memory and theory of mind as highly specialized. Additionally, we found support for guiding principles of brain organization generally, such that within-network connections contributed most to a given network’s specialization followed by connections from other specialized networks. These results have implications for identifying potential variations of network contributions in individuals with neurodevelopmental conditions.

## Introduction

Hemispheric specialization refers to a characteristic of brain organization in which specific functions draw on one hemisphere of the brain more than the other. These functional asymmetries give rise to reductions in redundancy (Levy, 1969), processing speed (Ringo et al., 1994), and interhemispheric conflict in function initiation (Andrew et al., 1982; Corballis, 1991). Importantly, disruptions to hemispheric specialization can have significant clinical implications, particularly in the context of neurodevelopmental and psychiatric conditions (X.-Z. Kong et al., 2022).

Measures of hemispheric specialization have previously ranged from the examination of split-brain patients (for review, see Gazzaniga, 2000) and brain lesions (Milner, 1971; Rasmussen & Milner, 1977) to the Wada test (Wada & Rasmussen, 1960) and intraoperative brain stimulation mapping (Penfield & Jasper, 1954). With the advent and development of functional neuroimaging, these methods now include many functional connectivity-based metrics. One such measure is the intrinsic laterality index (Liu et al., 2009), which quantifies within-versus between-hemisphere connectivity. Another includes the hemispheric contrast (Gotts et al., 2013), which examines node interactions between the two hemispheres through “segregation” (high within-hemisphere interactions) versus “integration” (high between-hemisphere interactions). Similarly, the autonomy index (Wang et al., 2014) captures a normalized ratio of within-versus between-hemisphere connectivity on the vertex level. Of these measures, the autonomy index holds particular interest since each vertex is taken as a region-of-interest, avoiding the influence of anatomical asymmetries.

Using the autonomy index, Wang et al. (2014) quantified specialization across seven functional networks and found that specialization was not restricted to a single left- or right-specialized network (Wang et al., 2014). Rather, the right frontoparietal control network and right ventral and dorsal attention networks, as well as the left default and frontoparietal control networks exhibited high degrees of specialization (see Fig. 5; Wang et al., 2014). The dual specialization of the frontoparietal control network evidences a joint coupling of executive control functions with a distinct pattern of networks in either hemisphere (Wang et al., 2014). While this and other studies have made significant contributions, much of what is known regarding hemispheric specialization has been derived from group-level analyses, an approach which is increasingly being exchanged for a within-individual “precision functional mapping” approach (Braga & Buckner, 2017; Gordon et al., 2017; Laumann et al., 2015).

In line with the precision neuroimaging approach and previous efforts to understand brain network organization and specialization, the present study examined two open questions. First, we explored which networks exhibit the greatest hemispheric specialization. Previously, Wang et al. (2014), identified networks associated with language, visuospatial attention, and executive control as being the most specialized. However, it remains unclear how these estimates might change with a greater number of examined networks, and when implemented at an individual level. In line with previous work, we hypothesized that networks associated with language, visuospatial attention, and executive control would show the greatest specialization. Second, we investigated which connections support specialization in a given network. Although a data-driven approach was implemented to address contributions to network specialization, we anticipated that the pattern of network contributions would follow a within-between network gradient or a specialization gradient. The within-between gradient hypothesis proposes that the connections contributing the most to a network’s specialization are those originating within the same network, as opposed to those from different networks. For example, connections between different areas within the language network would play a greater role in the specialization of the language network compared to connections between the language network and a different network, such as a visual network. The second hypothesis or specialization gradient hypothesis suggests that the connections contributing the most to a network’s specialization originate from other specialized networks as opposed to non-specialized networks. Under this hypothesis, one would expect that connections from a visuospatial attention or frontoparietal control network (i.e., specialized networks) would have a greater impact on the specialization of the language network than connections from a visual or somatomotor network (i.e., non-specialized networks).

## Materials and Methods

### Datasets and Overview

Three independent datasets were used for these analyses: The Human Connectome Project (HCP; split into discovery and replication datasets), the Human Connectome Project-Development (HCPD; Somerville et al., 2018), and the Natural Scenes Dataset (NSD; Allen et al., 2022). Each dataset was selected for its relatively high quantity of low-motion data per participant. See Peterson et al. (2023) for dataset descriptions and accompanying MRI acquisition parameters.

### MRI Processing

Processing for BOLD NIFTI files was comprised of the following steps: preprocessing, generation of individual parcellations, implementation of the autonomy index and the deconstructed autonomy index.

#### Preprocessing

Preprocessing took place on raw NIFTI files for the resting-state fMRI and task fMRI runs using a pipeline developed by the Computational Brain Imaging Group (CBIG; Kong et al., 2019; Li et al., 2019). The implementation of this pipeline was described previously (see Peterson et al., 2023) and is summarized briefly here. Following FreeSurfer surface reconstruction (FreeSurfer 6.0.1, RRID:SCR_001847; Dale et al., 1999), the pipeline includes the removal of the first four frames and motion correction (using FSL, RRID:SCR_002823; (Jenkinson et al., 2002; Smith et al., 2004), functional and structural image alignment (using FreeSurfer’s FsFast; Greve & Fischl, 2009), linear regression using multiple nuisance regressors (using a combination of CBIG in-house scripts and FSL MCFLIRT; Jenkinson et al., 2002), bandpass filtering (using CBIG in-house scripts), surface projection (using FreeSurfer’s *mri-vol2surf* function), and surface smoothing using a 6 mm full-width half-maximum kernel (using FreeSurfer’s *mri_surf2surf* function; Fischl et al., 1999).

#### Individual Network Parcellations

Following preprocessing, network parcellations were computed using a multi-session hierarchical Bayesian modeling (MS-HBM) pipeline (Kong et al., 2019) in MATLAB R2018b (RRID:SCR_001622; MATLAB, 2018). This pipeline generates parcellations for individuals with multiple sessions of fMRI data by using a variational Bayes expectation-maximization algorithm to learn group-level priors from a training dataset and apply those to estimate individual-specific parcellations. The model estimates various parameters, including group-level network connectivity profiles, inter-subject functional connectivity variability, intra-subject functional connectivity variability, a spatial smoothness prior, and an inter-subject spatial variability prior. The number of clusters (*k*) for all participants was set at 17 (Yeo et al., 2011).

Following the generation of individual parcellations, a Hungarian matching algorithm was used to match the clusters with the Yeo et al. (2011) 17-network group parcellation.

#### Autonomy Index

The autonomy index approaches specialization from a functional connectivity perspective and is known to reliably estimate specialization across neurotypical and clinical samples (Mueller et al., 2015; Sun et al., 2022; Wang et al., 2014). First, individual functional connectivity matrices were calculated for each BOLD run and then averaged across runs within an individual at the fsaverage6 resolution (40,962 vertices per hemisphere) in MATLAB R2018b (RRID:SCR_001622; MATLAB, 2018). From here, the autonomy index was computed. In summary, for each seed vertex obtained from a functional connectivity matrix, the degree of within-hemisphere connectivity and cross-hemisphere connectivity were computed by summing the number of vertices correlated to the seed in the ipsilateral hemisphere and in the contralateral hemisphere. This is then normalized by the total number of vertices in the corresponding hemisphere, thus the accounting for a potential brain size asymmetry between the two hemispheres. Finally, AI is calculated as the difference between normalized within- and cross-hemisphere connectivity as follows:

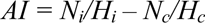

where *N_i_* and *N_c_* are the number of vertices correlated to the seed ROI (using a threshold of |0.25|) in the ipsilateral hemisphere and contralateral hemisphere, respectively. *H_i_* and *H_c_* are the total number of vertices in the ipsilateral and contralateral hemisphere, respectively. To compute the specialization of each functional network, the AI was averaged within the boundary of each network separately within each hemisphere on an individual basis, and then multiplied by 100.

Greater positive AI values indicate a higher ratio of within-hemisphere connections to between-hemisphere connections and are interpreted as greater network specialization.

#### Deconstructed Autonomy Index

The autonomy index serves as a general measure of specialization, and as such, it does not consider the specific networks responsible for contributing to the specialization of an individual network. In order to parse network specialization and address the aim of identifying contributions to network specialization, we formulated a deconstructed version of the autonomy index. This was accomplished by first calculating an average functional connectivity matrix for each individual as previously described. Then, for each target network (1-17) and each seed vertex derived from the average functional connectivity matrix, the degree of within- and cross-hemisphere connectivity was computed by summing the number of highly correlated vertices belonging to that target network in each hemisphere. This is normalized by the total number of vertices with a given target network label in each hemisphere (see Figure 1). This deconstructed AI (dAI) is calculated as follows:

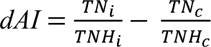

**Figure 1.**
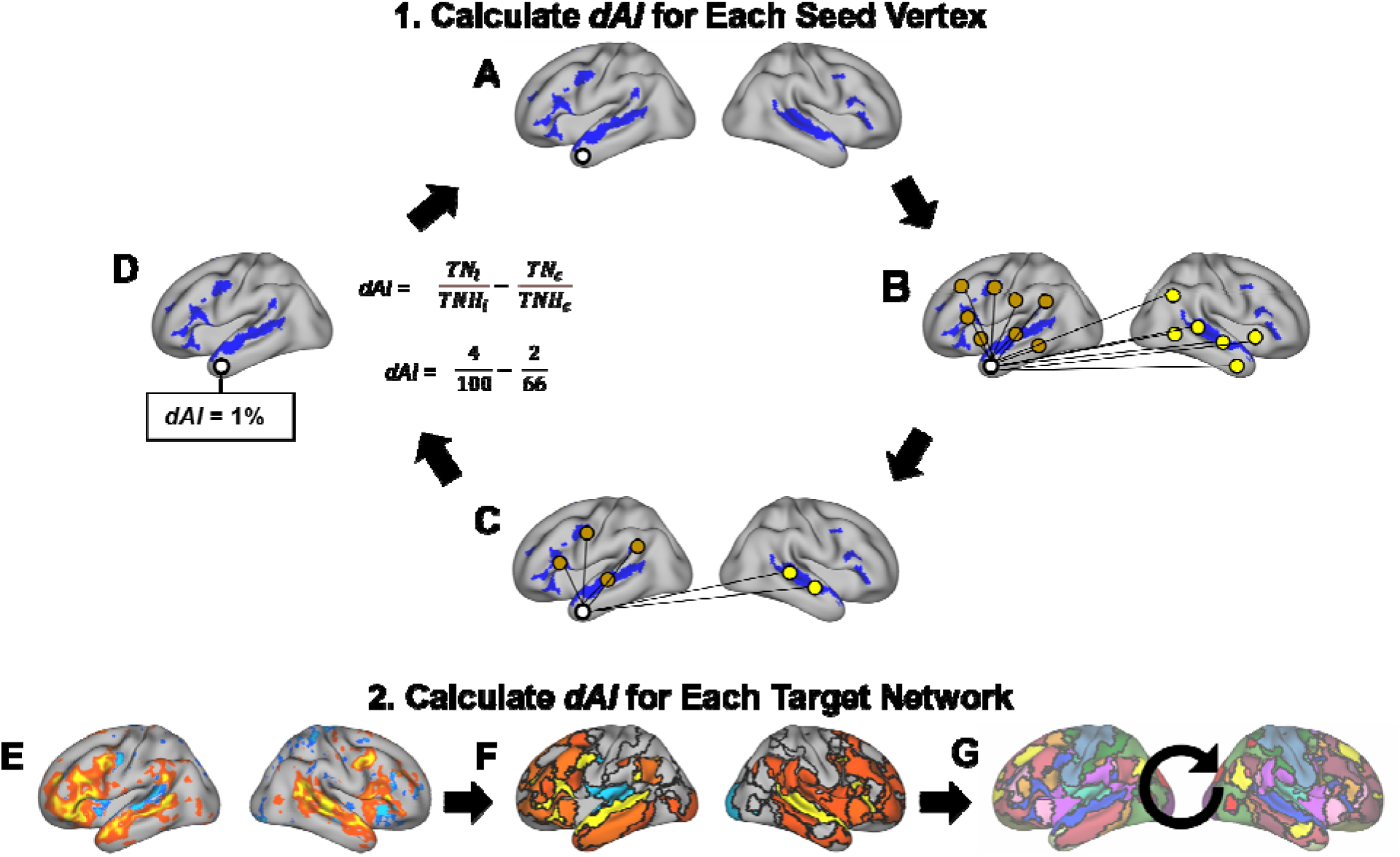
The deconstructed autonomy index. The first step consists of calculating the dAI for each vertex. Panel A depicts a seed vertex (white circle) and a target network (highlighted in blue). In Panel B, vertices correlated with the seed vertex (white circle) in the ipsilateral (brown circles) and contralateral (yellow circles) hemispheres are identified. In Panel C, the correlated vertices (brown and yellow circles) are filtered to those that fall within the boundaries of the target network. In Panel D, the dAI is calculated for the seed vertex (white circle), following the presented formula (see the main text for a description of the formula). In this example, the denominators of 100 and 66 are toy numbers and not representative of actual totals for network vertices. This process (Panels A-D) is repeated for each vertex, after which dAI is calculated for each target network. Panel E depicts the dAI values when language is the target network. Next, in Panel F, dAI values are averaged within each network and then multiplied by 100. This process is repeated for each target network (1-17; Panel G).

where *TN_i_* represents the number of vertices correlated with the seed vertex (using a threshold of |0.25|) that fall within the target network in the ipsilateral hemisphere, *TNH_i_* represents the number vertices with the target network label in the ipsilateral hemisphere, *TN_c_* represents the number of vertices correlated with the seed vertex that fall within the target network in the contralateral hemisphere, and *TNH_c_* represents the number of vertices with a given target network label in the contralateral hemisphere. This results in a matrix of dAI values for each target network for each subject. Then, for each target network matrix, the deconstructed AI is averaged within the boundaries of each network (1-17) and then multiplied by 100.

### Experimental Design and Statistical Analyses

The design of this study is largely within-individual, as will be detailed in the following sections. Analyses comprised a replication of Wang et al. (2014), reliability analyses (including test-retest reliability and a task effects analysis), the identification of specialized networks, and an analysis of within-network contributions. Statistical analyses took place in R 4.2.0 (RRID:SCR_001905; R Core Team, 2022).

### Replications of Wang et al. (2014)

Before expanding on the work of Wang et al. (2014), which was originally performed in Brain Genomics Superstruct Project subjects (Holmes et al., 2015), we first performed a replication in the HCP-Discovery and HCP-Replication datasets. This was accomplished by averaging left hemisphere and right hemisphere autonomy index values within the network boundaries of a seven-network group parcellation produced from 1000 subjects (the parcellation is freely available online: https://surfer.nmr.mgh.harvard.edu/fswiki/CorticalParcellation_Yeo2011; Yeo et al., 2011).

Then, to proceed in a step-wise fashion, the same procedure was undertaken with a 17-network group parcellation (Yeo et al., 2011).

#### Reliability Analyses

Reliability analyses sought to address the following questions: 1) What is the test-retest reliability of the autonomy index using individual parcellations, and 2) Is there a task effect on autonomy index estimation?

##### Test-Retest Reliability

In order to determine the test-retest reliability of the autonomy index, HCP subjects with all four resting-state runs available after preprocessing were utilized (*N* = 232). Individual parcellations were generated separately for the first two runs (the first scanning session) and the second two runs (the second scanning session). Functional connectivity matrices were also generated separately for the first session and the second session, and the autonomy index was calculated on both. Next, the autonomy index was averaged within network boundaries for the left and right hemispheres for each session. Outliers were fenced on a network basis to an upper limit of the third quartile plus 1.5 multiplied by the interquartile range, and a lower limit of the first quartile minus 1.5 multiplied by the interquartile range. Finally, an intraclass correlation was calculated for the averaged autonomy index values for the top five most left-specialized(language, dorsal attention-A, default-A, default-C, and limbic-B) and the top five most right-specialized networks (salience/ventral attention-A, control-B, control-C, default-C, and limbic-B). Intraclass correlations were then evaluated using the standard guidelines from Koo & Li (2016), with values less than 0.5 indicating poor reliability, values between 0.5 and 0.75 indicating moderate reliability, values between 0.75 and 0.9 indicating good reliability, and values greater than 0.9 indicating excellent reliability (based on a 95% confidence interval). Spearman rank correlations were then used to examine potential relationships between network test-retest reliability and a network-averaged signal-to-noise ratio.

##### Task Effects on the Autonomy Index

Following the procedure outlined in Peterson et al. (2023), task effects were also examined for the estimation of the autonomy index within individuals. Briefly, individual parcellations were generated using both task and resting-state fMRI data from the NSD dataset using various combinations of runs within task type: even-numbered runs, odd-numbered runs, the first half of runs, the second half of runs, and two random selections of runs (without replacement). Intraclass correlation coefficients were then used to compare autonomy index overlap within task and between tasks for each hemisphere. Wilcoxon Signed Rank tests were then used to compare the intraclass correlations (R Core Team, 2011; Wilcoxon, 1945).

#### Identifying Specialized Networks with Individual Parcellations

After establishing the reliability of approaching specialization from an individual-level perspective using the autonomy index, we addressed the first hypothesis of determining whether any of the 17 networks exhibited specialization, and if so, which exhibited the greatest specialization. The following analyses were first implemented in the HCP-Discovery dataset and then replicated in the HCP-Replication and HCPD datasets using all data available from each participant. First, to determine whether any networks exhibited specialization, multiple regressions were implemented for each of the 17 networks, separately for the left and right hemispheres. Models consisted of a given network’s mean autonomy index value and the covariates of mean-centered age, sex, mean-centered mean framewise displacement, and handedness (measured via the Edinburgh Handedness Inventory; Oldfield, 1971). A network was considered specialized if the model intercept was significant at the Bonferroni-corrected alpha level of 0.001. The top five left- and right-lateralized networks were determined using the HCP-Discovery dataset.

#### Within-Network Contributions

To address our second hypothesis regarding contributions to specialized networks, we implemented the deconstructed autonomy index in the top five left- and right-specialized networks as determined via the HCP-Discovery dataset. These values were adjusted for mean-centered age, mean-centered mean framewise displacement, sex, and handedness. Following model-adjustment, potential patterns were visually identified and then assessed quantitatively via matrices of mean dAI values first within the HCP-Discovery dataset and then within the HCP-Replication and HCPD datasets.

#### Code Accessibility Statement

With the exception of the HCPD dataset, the data reported on in the present study can be accessed publicly online (HCP: https://db.humanconnectome.org/; NSD: http://naturalscenesdataset.org/). The HCPD dataset is hosted through the NIMH Data Archive (NDA) through which access may be requested. Preprocessing and individual parcellation pipeline code are available through the CBIG repository on GitHub at https://github.com/ThomasYeoLab/CBIG. Scripts used to implement the processing pipelines and perform statistical analyses are also available on GitHub at https://github.com/Nielsen-Brain-and-Behavior-Lab/AutonomyIndex2023.

## Results

### Replication of Wang et al. (2014)

Using the Yeo et al. (2011) seven-network group parcellation, we identified the default and frontoparietal control networks as the most left-specialized networks, and the frontoparietal control, dorsal attention and ventral attention networks as the most right-specialized networks for both the HCP-Discovery and HCP-Replication datasets (see Supplementary Figure S1). This pattern of specialized networks replicates that found by Wang et al. (2014). Next, we expanded on the Wang et al. (2014) analysis to examine network specialization using the Yeo et al. (2011) 17-network group parcellation. We identified the dorsal attention-A, language, default-A, default-C, and limbic-B networks as the most left-specialized across both datasets (see Supplementary Figure S2). The control-B, default-C, and limbic-B networks showed the greatest right-hemisphere specialization.

#### Reliability Analyses

Following the group-based parcellation analyses, the autonomy index was adapted to the individual through individual network parcellations. However, prior to examining specialization, the reliability of this individualized approach was examined through the following test-retest reliability and task effects analyses.

#### Test-Retest Reliability

Using HCP subjects with all four resting-state runs available after preprocessing (*N* = 232), test-retest reliability was assessed for the top five left- and right-specialized networks. For the left-specialized networks, intraclass correlations were within a moderate range, between 0.55 to 0.77, with the lowest being the limbic-B network (ICC = 0.55, *F*(231, 231) = 3.5, *p* < .001, 95% CI [0.46, 0.63]; see Figure 2). For the right-specialized networks, the intraclass correlations were also in the moderate range, from 0.55 to 0.72, with the control-C network exhibiting the lowest reliability (ICC = 0.55, *F*(231, 231) = 3.5, *p* < .001, 95% CI [0.45, 0.63]). Spearman rank correlations identified no relationship between test-retest reliability (intraclass correlation coefficients) and network-averaged temporal signal-to-noise ratios (left hemisphere: *r*(15) = 0.11, *p* = .68; right hemisphere: *r*(15) = .27, *p* = .29; see Supplementary Figure S3).

**Figure 2.**
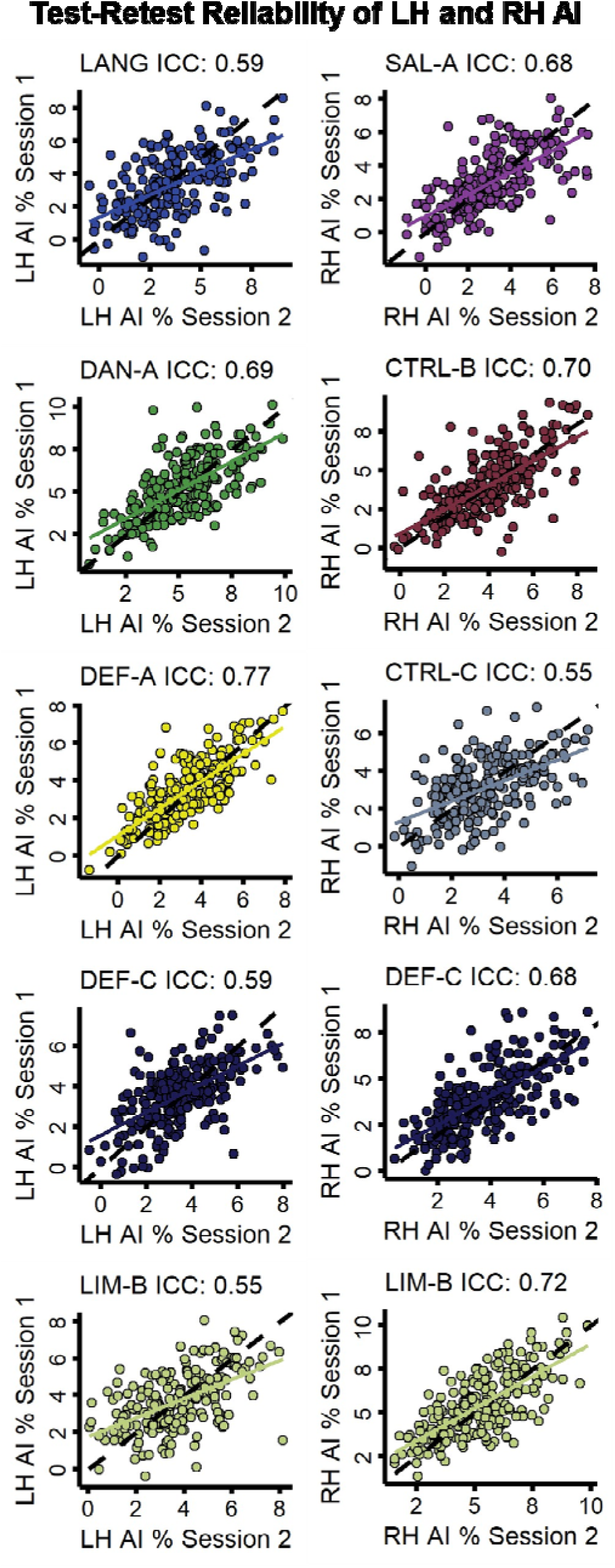
Test-retest reliability of autonomy index values for five left- and right-specialized networks in 232 HCP subjects. Left-specialized networks (left column) included language (LANG), dorsal attention-A (DAN-A), default-A (DEF-A), default-C (DEF-C), and limbic-B (LIM-B). Right-specialized networks (right column) included salience/ventral attention-A (SAL-A), control-B (CTRL-B), control-C (CTRL-C), default-C (DEF-C), and limbic-B (LIM-B). In each plot, a circle represents a subject, and the dashed identity line in black represents the theoretical perfect correspondence between the two sessions.

#### Task Effects on the Autonomy Index

Using the NSD dataset (*N* = 8) to compare potential differences between resting-state and task fMRI on autonomy index estimates for the left and right hemispheres, we found differences between the within-task comparisons and between task comparisons for autonomy index intraclass correlation coefficients (see Figure 3). Wilcoxon signed rank comparisons revealed a difference in within-task (Task-Task and Rest-Rest) intraclass correlation coefficients for even versus odd numbered runs (LH: *V* = 33, *p* = .04; RH: *V* = 34, *p* = .02) as well as for the first half versus the second half of runs (LH: *V* = 34, *p* = .02; RH: *V* = 32, *p* = .05), but not for the random selection of runs (LH: *V* = 26, *p* = .31; RH: *V* = 24, *p* = .46). Regardless of how the data were split, a task effect in intraclass correlation coefficients was found between within-task (Task-Task) and between-task (Task-Rest) intraclass correlation coefficients for even versus odd numbered runs (LH: *V* = 36, *p* = .008; RH: *V* = 35, *p* = .02), the first half versus the second half of runs (LH: *V* = 36, *p* = .008; RH: *V* = 36, *p* = .008), and the random selection of runs (LH: *V* = 34, *p* = .02; RH: *V* = 36, *p* = .008).

**Figure 3.**
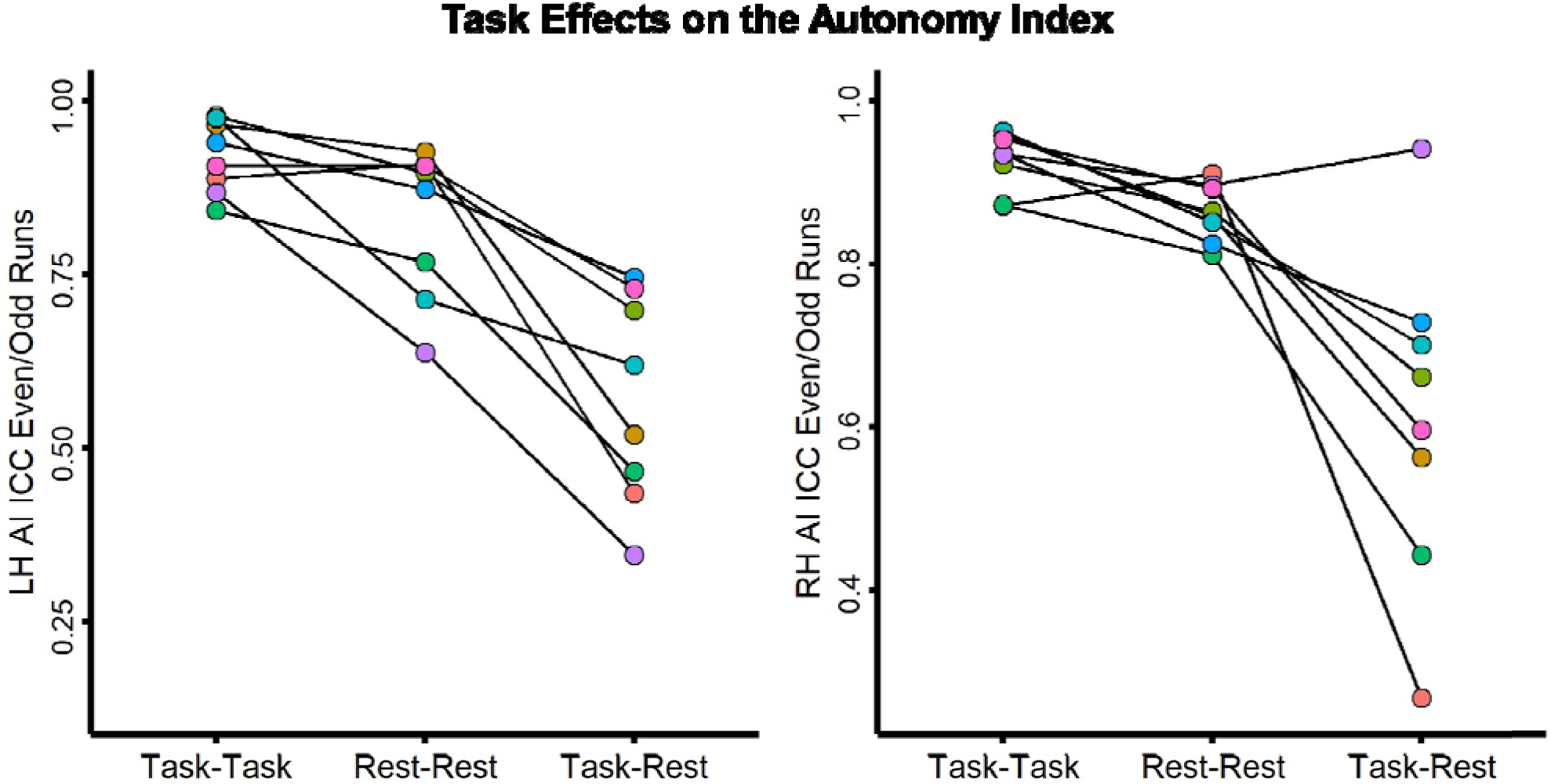
Task effects on the autonomy index in the NSD dataset. In the figure, intraclass correlation coefficients for the left and right hemisphere autonomy indices are shown for each participant, comparing even-versus odd-numbered runs (with the left hemisphere values shown on the left and right hemisphere values shown on the right). Regardless of how the data were split (even-versus odd-numbered runs, the first half versus the second half, or a random selection without replacement), a task effect was found.

### Identifying Specialized Networks with Individual Parcellations

Following the investigations into the reliability of the autonomy index as implemented in individuals, we addressed the major hypotheses regarding which networks were specialized and which connections contributed to this specialization. A series of multiple regressions were used to identify if any of the 17 networks were specialized, first in the HCP-Discovery dataset and then in the HCP-Replication and HCPD datasets. Networks with significant left-hemisphere specialization (*p* < .001) across all three datasets included each network except visual-A and default-B (see Supplementary Table 1). All 17 networks were found to have significant right-hemisphere specialization (*p* < .001) across all three datasets (see Supplementary Table 2). Of the covariates, only handedness was reliably significant across all three datasets for the left-hemisphere averaged salience/ventral attention-A autonomy index (see Supplementary Figure S4). See Figure 4 for model-adjusted mean autonomy index values for each of the 17 networks for the left hemisphere (Panel A) and the right hemisphere (Panel B). Notably, the limbic-B and default-C networks appear to be strongly specialized to both hemispheres, similar to what has previously been observed with the frontoparietal control network by Wang and colleagues (Wang et al., 2014). The most specialized networks were identified as the top five left- and right-lateralized networks from the HCP-Discovery dataset.

**Figure 4.**
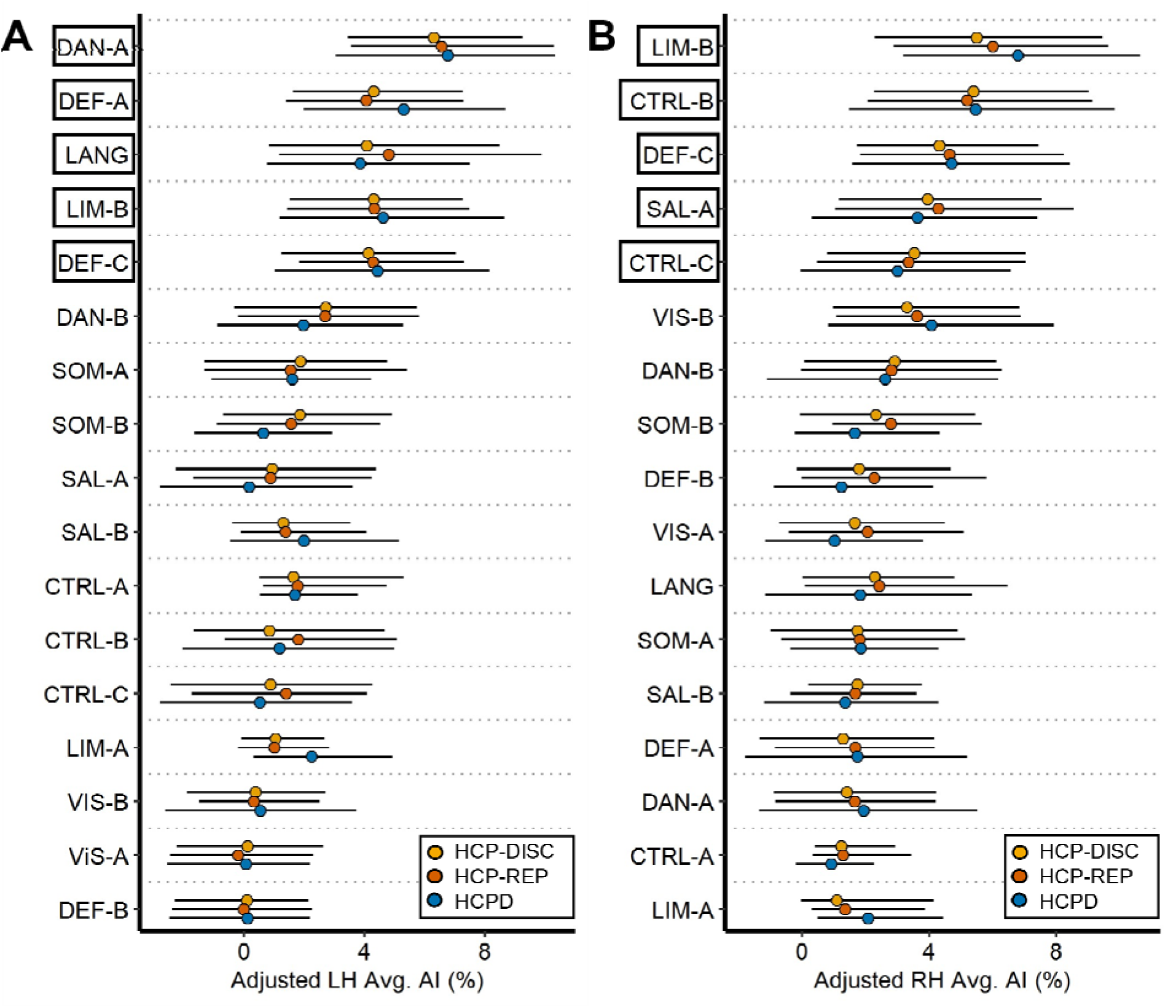
Specialization for 17 networks across the HCP-Discovery, HCP-Replication, and HCPD datasets. For each panel, the y-axis displays the 17 networks and the x-axis displays the adjusted average autonomy index values, with greater values representing greater hemispheric specialization (left hemisphere on Panel A and right hemisphere on Panel B). Autonomy index values were adjusted by regressing out the effects of mean-centered age, mean-centered mean framewise displacement, sex, and handedness using the following formula: AI_adj_ = AI_nat_ — [β_1_(mean-centered age_nat_ – mean of mean-centered age_nat_) + β_2_(mean-centered FD_nat_ – mean of mean-centered FD_nat_) + β_3_(sex_nat_ – mean sex_nat_) + β_4_(handedness_nat_ – mean handedness_nat_)]. Autonomy index adjustment occurred separately for each network within each dataset for each hemisphere. Bars represent the 2.5 and 97.5 percentiles, and black boxes have been placed around the top five left- and right-specialized networks (determined using the HCP-Discovery dataset).

### Comparison of Group and Individualized Approaches

A key assumption of the present study has been that an individualized approach to network specialization would elicit a more precise estimate of the autonomy index than group-based approaches, an assumption backed by evidence demonstrating advances arising from a precision approach to neuroimaging (Braga et al., 2020; Braga & Buckner, 2017; DiNicola et al., 2020; Gordon et al., 2017; Gratton et al., 2018; Laumann et al., 2015).Visual comparison of the networks with the greatest left- and right-hemisphere specialization using a group 17-network parcellation (Supplementary Figure S2) and 17-network individual parcellations (Figure 4) reveals nearly identical results. This challenges our initial assumption and implies that, in the context of the present study, the choice of network parcellation method may have a limited impact on the estimation of the autonomy index.

### Within-Network Contributions to Specialization

Next, to address the second hypothesis and decompose network specialization to identify the greatest contributions to each network’s specialization, deconstructed autonomy index values were averaged within the 17 networks for each target network. As described in the Methods section, the dAI is calculated as a ratio of within- and between-hemisphere connectedness for each vertex and target network. Visual examination of the top left- and right-specialized networks (determined using the HCP-Discovery dataset) initially indicated that within-network connections appear to be the greatest contributors. For example, language network connections contribute the most to language network specialization (see Supplementary Figures S5-S10).

Following within-network connections, other specialized networks appear to be the second largest contributor to specialization (see Figure 5 and Supplementary Figure S11). As an example of this second pattern, it is more likely that the specialized default-C network is contributing to the specialization of the language network than a network that isn’t specialized, such as visual-A. Matrices of mean dAI scores for all potential 17 target networks confirmed these two principles (see Supplementary Figures S12-S14).

**Figure 5.**
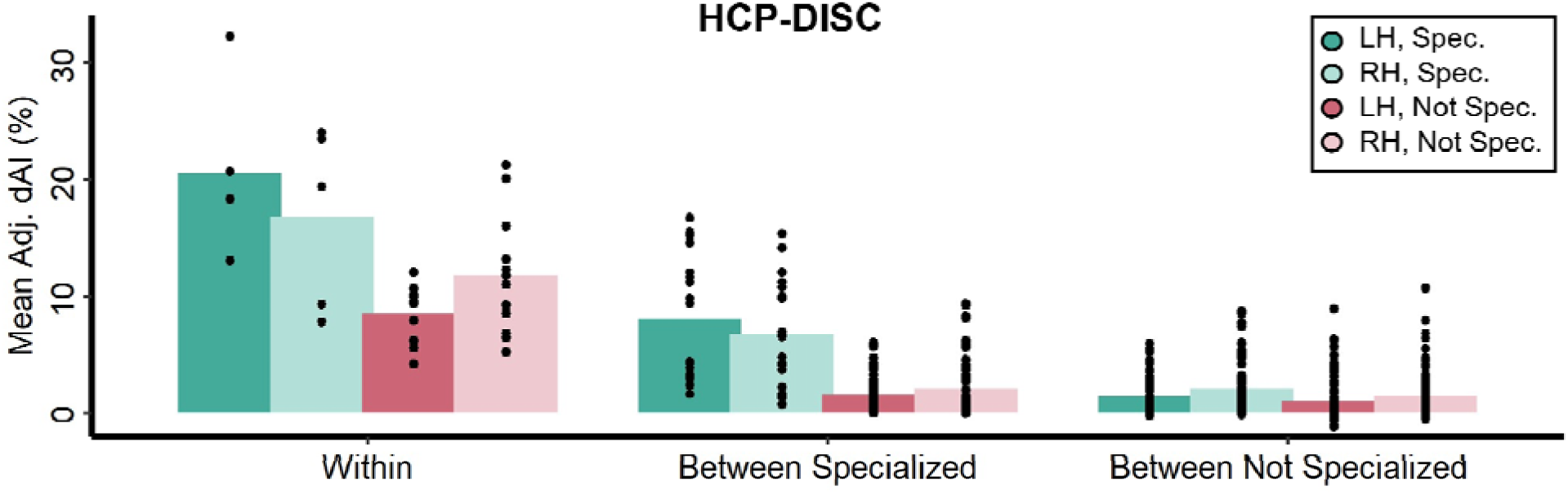
Deconstructed autonomy index (dAI) values within and between specialized networks for the HCP-Discovery dataset. dAI scores were grouped as being within-network (e.g., target network language and averaged network language), between specialized networks (e.g., target network language and averaged network dorsal attention-A), and between not specialized (e.g., target network language and averaged network visual-A). Specialized networks are the top five left- and right-specialized networks (indicated with black boxes in Figure 4). Next, dAI scores were further binned depending on the target network as being specialized or not specialized (specialized networks included the top five left- and right-specialized networks). Finally, dAI scores were organized by hemisphere, left or right. Each point represents a single target and averaged network combination mean dAI score. Across each dataset, within-network contributions appear to be greatest followed by between-specialized network contributions and then between not-specialized network contributions (see Supplementary Figure S11 for the HCP-Replication and HCPD datasets).

## Discussion

In the present study, we examined brain network specialization using a functional connectivity-based measure, replicating and expanding on the work of Wang et al. (2014). After directly replicating Wang et al. (2014) and addressing the reliability of a within-individual implementation, we identified specialized networks at a 17-network resolution and determined the greatest contributions to specialization. In line with prior work, we identified a highly left-specialized network supporting language function and several highly right-specialized networks supporting visuospatial attention and executive control functions (Wang et al., 2014). However, we identified networks other than those directly associated with language, visuospatial attention, and executive control as being the most specialized. Additionally, across three datasets, we found that within-network connections contributed most to a given network’s specialization followed by connections from other specialized networks. Taken as a whole, these results provide evidence for guiding principles of brain organization generally and the specialization of macroscale brain networks specifically.

### Evidence for the Reliability of the Autonomy Index

In the present study, we examined specialization using the autonomy index. However, the implementation of this measure differed from the original in two key ways: the use of a 17-network parcellation scheme as opposed to seven networks, and the adoption of individual parcellations to delineate network boundaries as opposed to a single group-level network parcellation. While we contend that these differences facilitated the acquisition of novel insights into network specialization, it was uncertain as to whether they compromised the integrity of the autonomy index. However, visual comparison between the most specialized networks resulting from the 17-network individual and group parcellations reveals nearly identical results. This finding contradicts our assumption regarding the optimality of individual parcellations and suggests that within the scope of this study, the use of individual network parcellations exerted minimal influence on the calculation of the autonomy index. Additionally, when using the individual parcellations, the top five left- and right-specialized networks fell within moderate test-retest reliability. This also indicates that the use of individual parcellations did not adversely affect the detection of signal and thus specialization. In other words, if we were measuring noise, we would expect very low test-retest reliability.

While the individual-level implementation of the autonomy index was found to be reliable, task effects were found. This finding replicates previously observed task effects on a different measure of functional connectivity-derived network specialization (see Peterson et al., 2023). This is likely due to the idea that the act of “resting” introduces greater variability in functional connectivity compared to that observed during task-based fMRI, potentially as a result of mind wandering (Elton & Gao, 2015; Finn & Bandettini, 2021). Furthermore, when it comes to predicting individual traits, task-based models outperform rest-based models, and this has been attributed to the “unconstrained nature” of the resting state (Greene et al., 2018).

### Identification of Highly Specialized Networks

Notably, the top five greatest left-specialized networks included the language, dorsal attention-A, default-A, default-C, and limbic-B networks, while the right-specialized counterparts included the salience/ventral attention-A, control-B, control-C, default-C, and limbic-B networks. Similar to previous work, a highly left-specialized network supports language function (such as the language network) and several highly right-specialized networks support visuospatial attention and executive control functions (such as the salience/ventral attention-A, control-B, and control-C networks; Wang et al., 2014). While networks associated with language, attention, and executive control were highly specialized, they were not the most specialized left- and right-specialized networks. Instead, the dorsal attention-A and limbic-B networks were the most left- and right-specialized networks, respectively. Corroborating this finding, a network surface area-based approach to specialization also using the Kong et al. (2019) 17-network individual parcellation identified these two networks as the most specialized or lateralized (see Peterson et al., 2023). While it is unclear why the language and visuospatial attention networks were not identified as the most specialized networks, this finding may be due to the larger number of networks examined here, compared with the seven networks examined by Wang et al. (2014). Regardless, this result was robust across the 17-network group-averaged parcellation and the individual parcellations, indicating that the method of parcellation (group or individual) had little influence on this result.

An alternative explanation for the identification of the dorsal attention-A and limbic-B networks as being most specialized comes from other within-individual investigations. Work on the default network found that the group-defined default network is split into two parallel yet interdigitated networks which subserve different functions on the individual level (Braga & Buckner, 2017; DiNicola et al., 2020). Thus, it may be that, by taking a fine-grained approach to network parcellations, networks which have previously been described as bilateral may be split into subnetworks, of which one may be left- or right-specialized. In the case of dorsal attention, previously described as belonging to a bilateral network (Fox et al., 2006; Mengotti et al., 2020), this may have been split into a highly left-specialized network (dorsal attention-A) and a less specialized network (dorsal attention-B).

The present study did not replicate dually-specialized frontoparietal control networks (Wang et al., 2014) within the 17-network parcellation. Instead, we identified the control-B and control-C networks as being highly right-specialized, with minimal specialization of the control-A network. Additionally, we found that the default-C and limbic-B networks were highly specialized to both hemispheres. Further investigation is needed to show why these specific networks show this dual or coupling pattern of specialization.

### Contributions to Specialization

In order to better understand which functional connections contribute the most to a given network’s specialization, we implemented a novel version of the autonomy index: the deconstructed autonomy index. Resulting patterns of dAI values indicated that the greatest contributions to network specialization were from the same network (following a within-between network gradient) followed by other specialized networks (following a specialization gradient). This first principle of specialization contributions--that within-network contributions are greatest--is supported by the idea that hemispheric asymmetries increase the modularization of functions, thereby decreasing redundancy (Levy, 1969), processing speed (Ringo et al., 1994), and interhemispheric conflict (Andrew et al., 1982; Corballis, 1991; Gerrits et al., 2020; Vallortigara, 2006). A larger number of within-network connections would contribute to that network’s specialization of function and likely increase the efficiency of that network. Support for the second principle of network contributions—that connections from other specialized networks make up the second largest contribution—can be found from graph theoretical analysis and so-called “rich clubs”. These hubs are composed of high degree and high strength nodes, and are highly interconnected to one another (Colizza et al., 2006; Opsahl et al., 2008). Notably, rich club nodes have been identified in highly integrated brain regions, such as cingulate and pericingulate regions, as well as highly specialized brain regions including the orbitofrontal cortex, caudate, fusiform gyrus, and hippocampus (Kocher et al., 2015). Relevant to the present study, this work with rich club hubs evidences the following idea: specialized brain regions can be highly interconnected with one another. While the specialized brain networks discussed in the present study are not necessarily the same as the specialized brain regions referenced in Kocher et al. (2015), our work suggests that the strength of between-network contributions is less than within-network contributions.

### Limitations and Future Directions

In order to obtain individual parcellations that are reliable within individuals and comparable with previous work, the MS-HBM pipeline was selected. As a part of this pipeline, inter-subject variability is accounted for by way of group priors (derived from 37 Brain Genomics Superstruct Project subjects). One risk with incorporating group priors is that they may constrain network boundaries in ways that do not necessarily reflect an individual’s functional neurobiology. As noted in Kong et al. (2019), functional connectivity profiles derived from individuals are fairly noisy compared with those from a group-averaged profile. Thus, while the incorporation of group priors may reduce this noise, some signal may be lost as well.

An additional limitation with the selected method concerns the failure to capture the moment-to-moment dynamics of brain function. By creating static parcellations, we have oversimplified the temporal dimension of these large-scale networks.

Further applications of the deconstructed autonomy index could involve studying populations other than neurotypical children, adolescents, and adults. For example, it would be interesting to know how contributions to network specialization may be different in autism, for which atypical functional lateralization patterns have been reliably observed (Anderson et al., 2010; Cardinale et al., 2013; Eyler et al., 2012; Harris et al., 2006; Jouravlev et al., 2020; Kleinhans et al., 2008; X.-Z. Kong et al., 2022; Lindell & Hudry, 2013; Müller et al., 2003; Nielsen et al., 2014; Persichetti et al., 2022; Redcay & Courchesne, 2008) as well as schizophrenia (Agcaoglu et al., 2018; Ocklenburg et al., 2013; Sommer et al., 2001).

Additionally, it is unknown when the patterns of network contributions are established or how they potentially change in older adulthood. Future research could provide valuable insights into potential variations of network contributions in individuals with neurodevelopmental conditions and shed light on the developmental trajectory and potential changes in network specialization throughout adulthood.

## Conclusions

In the present study, we examined brain network organization on an individual level through the lens of specialization. We identified most networks as exhibiting greater within-hemisphere connectivity than between-hemisphere connectivity. Additionally, we found that the greatest contributors to network specialization were first within-network connections followed by connections with other specialized networks.

## Acknowledgements

Data were provided in part by the Human Connectome Project, WU-Minn Consortium (principal Investigators: David Van Essen and Kamil Ugurbil; 1U54MH091657) funded by the 16 NIH Institutes and Centers that support the NIH Blueprint for Neuroscience Research; and by the McDonnell Center for Systems Neuroscience at Washington University. HCPD data reported in this publication was supported by the National Institute of Mental Health of the National Institutes of Health under Award Number U01MH109589 and by funds provided by the McDonnell Center for Systems Neuroscience at Washington University in St. Louis. The HCP-Development 2.0 Release data used in this report came from DOI: 10.15154/1520708.

Collection of the NSD dataset was supported by NSF IIS-1822683 and NSF IIS-1822929. DLF is supported by funding from the European Union’s Horizon 2020 research and innovation programme under the Marie Skłodowska-Curie grant agreement No 101025785. Furthermore, we acknowledge the support of the Office of Research Computing at Brigham Young University.

## Author Contributions

M.P.: Conceptualization, Methodology, Software, Validation, Formal analysis, Investigation, Data curation, Writing – original draft, Writing – review & editing, Visualization, and Project administration. D.L.F.: Writing – review & editing. J.A.N.: Conceptualization, Methodology, Writing — review and editing, Supervision, and Project administration.

