## Supplemental Figures and Tables for "Parsing Brain Network Specialization: A Replication and Expansion of Wang et al. (2014)"

**A Replication and Expansion of Wang et al. (2014)**

Supplementary Materials

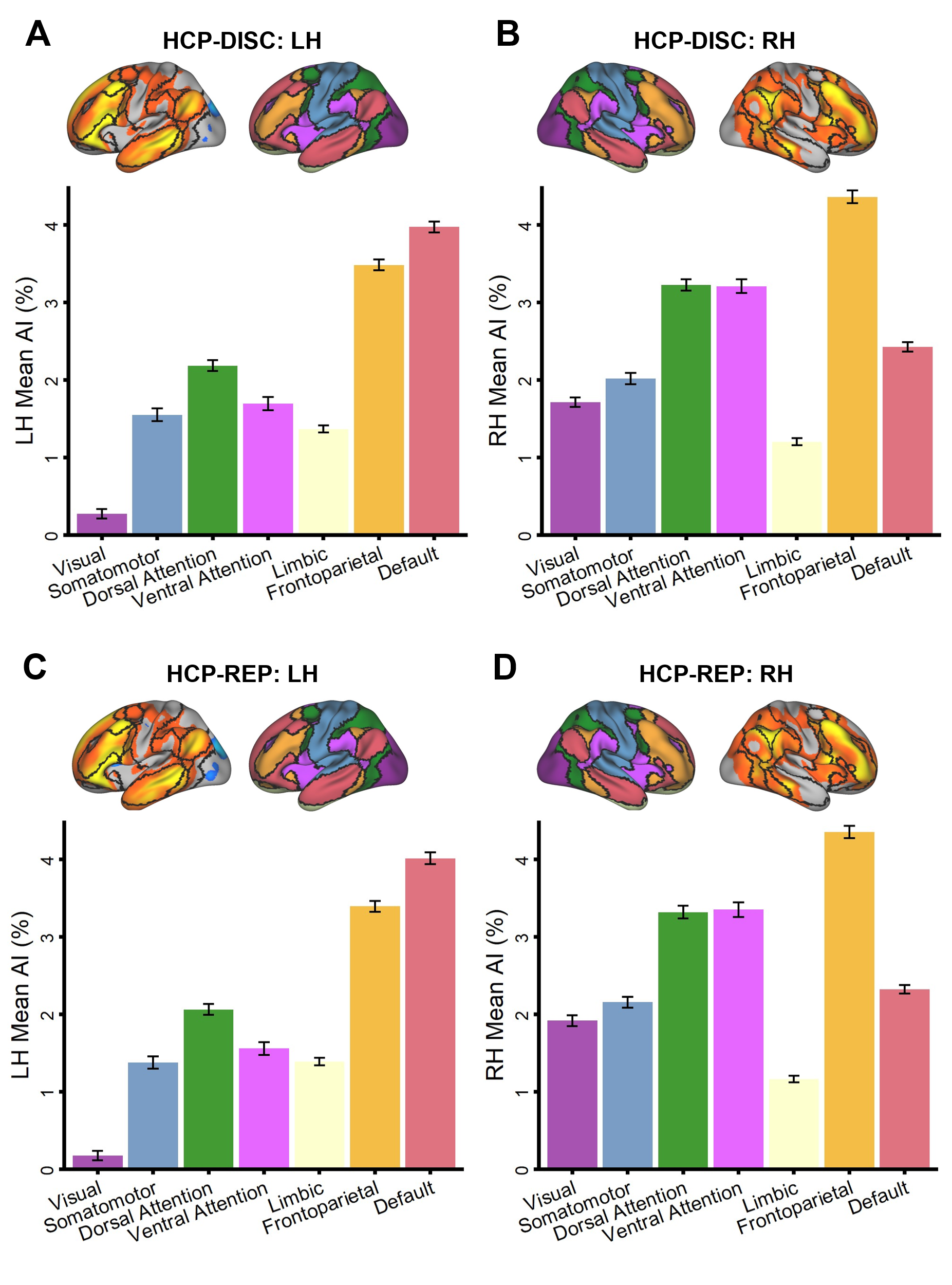
**Figure S1.** Direct replication of Wang et al. (2014) across the HCP-Discovery and HCP-Replication datasets. Panel A depicts autonomy index (AI) values averaged within the left hemisphere networks of the HCP-Discovery dataset. Panel B depicts AI values averaged within the right hemisphere networks of the HCP-Discovery dataset. Panel C depicts AI values averaged within the left hemisphere networks of the HCP-Replication dataset. Panel D depicts AI values averaged within the right hemisphere networks of the HCP-Replication dataset. Network boundaries came from a 1000-subject parcellation (Yeo et al., 2011). Error bars represent the standard error of the mean.

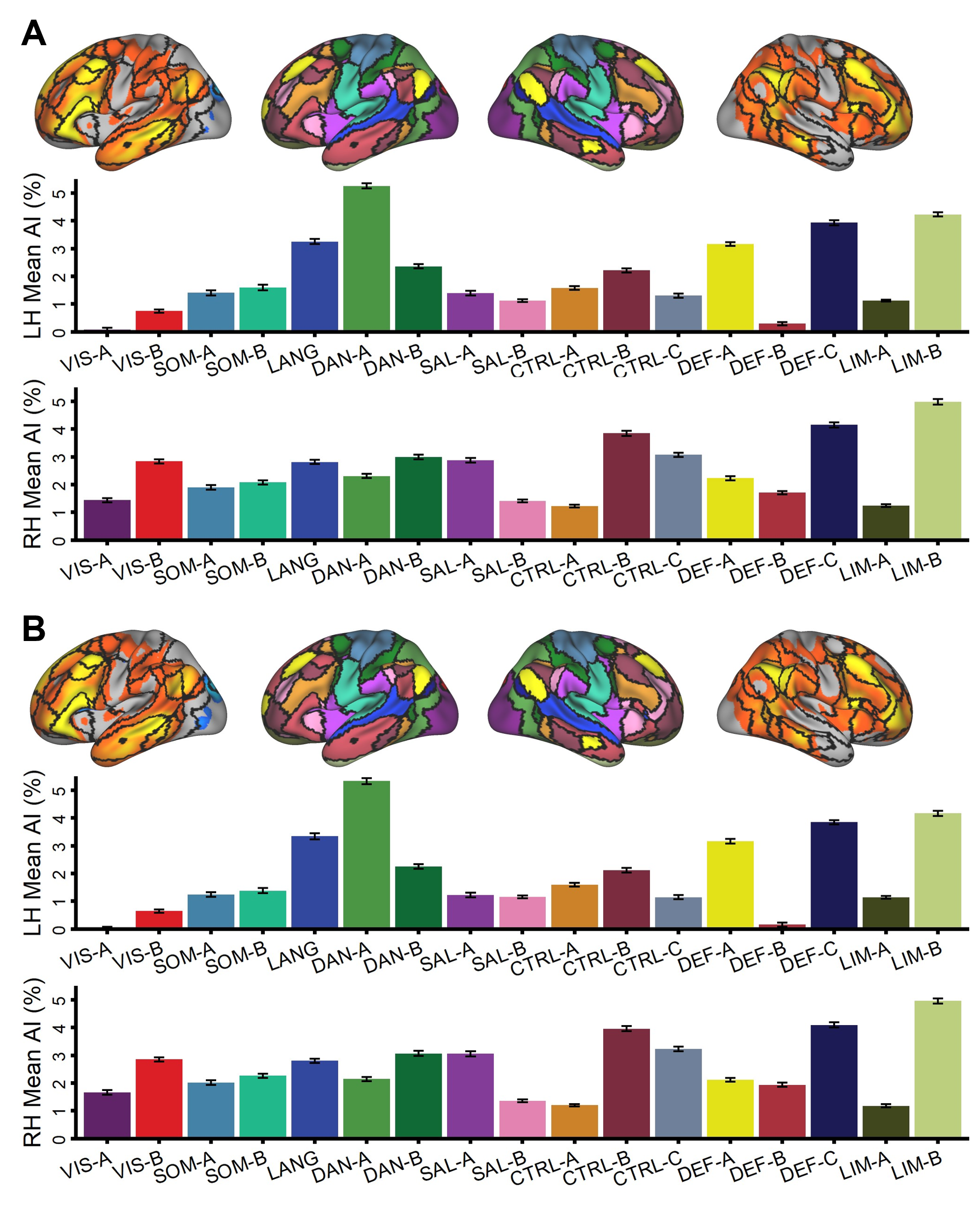
**Figure S2.** Replication of Wang et al. (2014) across 17 networks in the HCP-Discovery and HCP-Replication datasets. Panel A depicts the average autonomy index (AI) values for each of 17 networks in the left and right hemispheres for the HCP-Discovery dataset. Panel B depicts the average AI values for each of 17 networks in the left and right hemispheres for the HCP-Replication datasets. Network boundaries came from a 1000-subject parcellation (Yeo et al., 2011). Error bars represent the standard error of the mean.

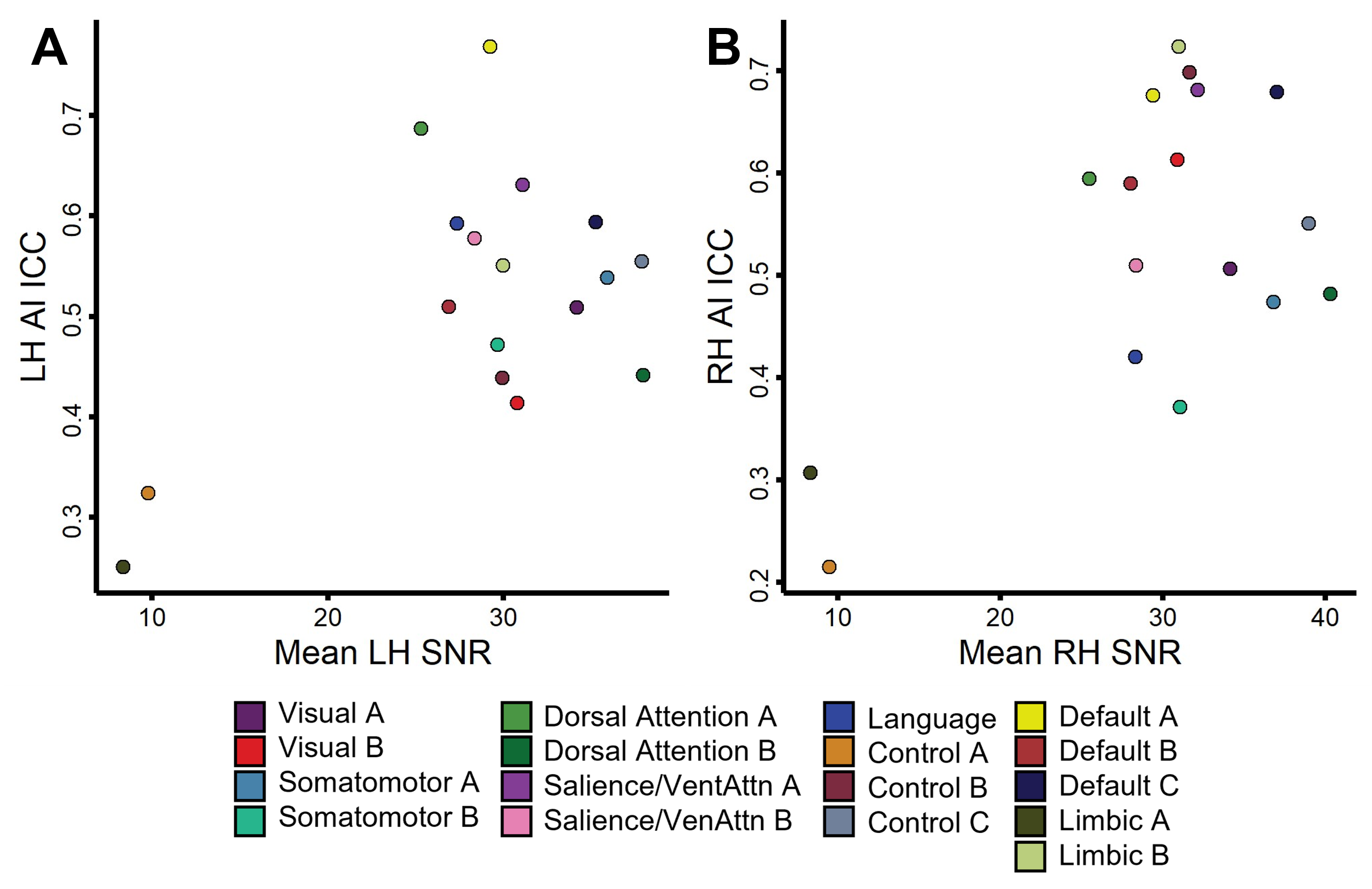

**Figure S3.** Test-retest reliability and temporal signal-to-noise ratios (tSNR) by network. tSNR was calculated by taking the mean BOLD signal and dividing it by the standard deviation in BOLD signal across all vertices for each participant. The tSNR was then averaged within network boundaries for each participant before being averaged across participants. Spearman rank correlations identified no relationship between intraclass correlation coefficients and network-averaged temporal signal-to-noise ratios in the left hemisphere (r(15) = 0.11, *p* = .68; Panel A) and right hemisphere (r(15) = .27, *p* = .29; Panel B). Each circle represents a single network.

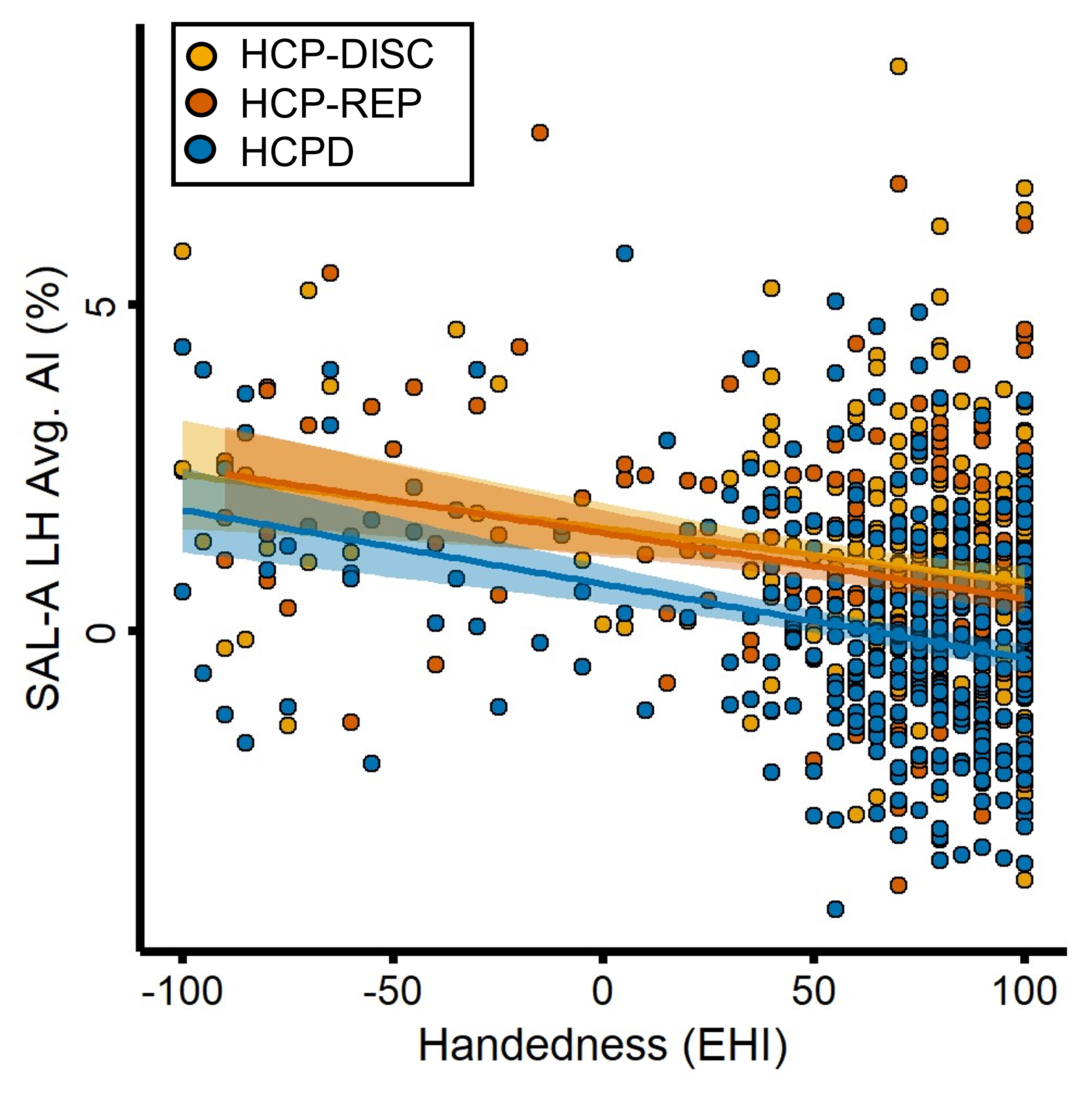
**Figure S4.** Negative relationship between handedness and salience/ventral attention-A left hemisphere mean autonomy index. Across the HCP-Discovery, HCP-Replication, and HCPD datasets, handedness (measured via the Edinburgh Handedness Inventory) was a significant covariate for the left hemisphere salience/ventral attention-A mean autonomy index. Each point represents a subject.

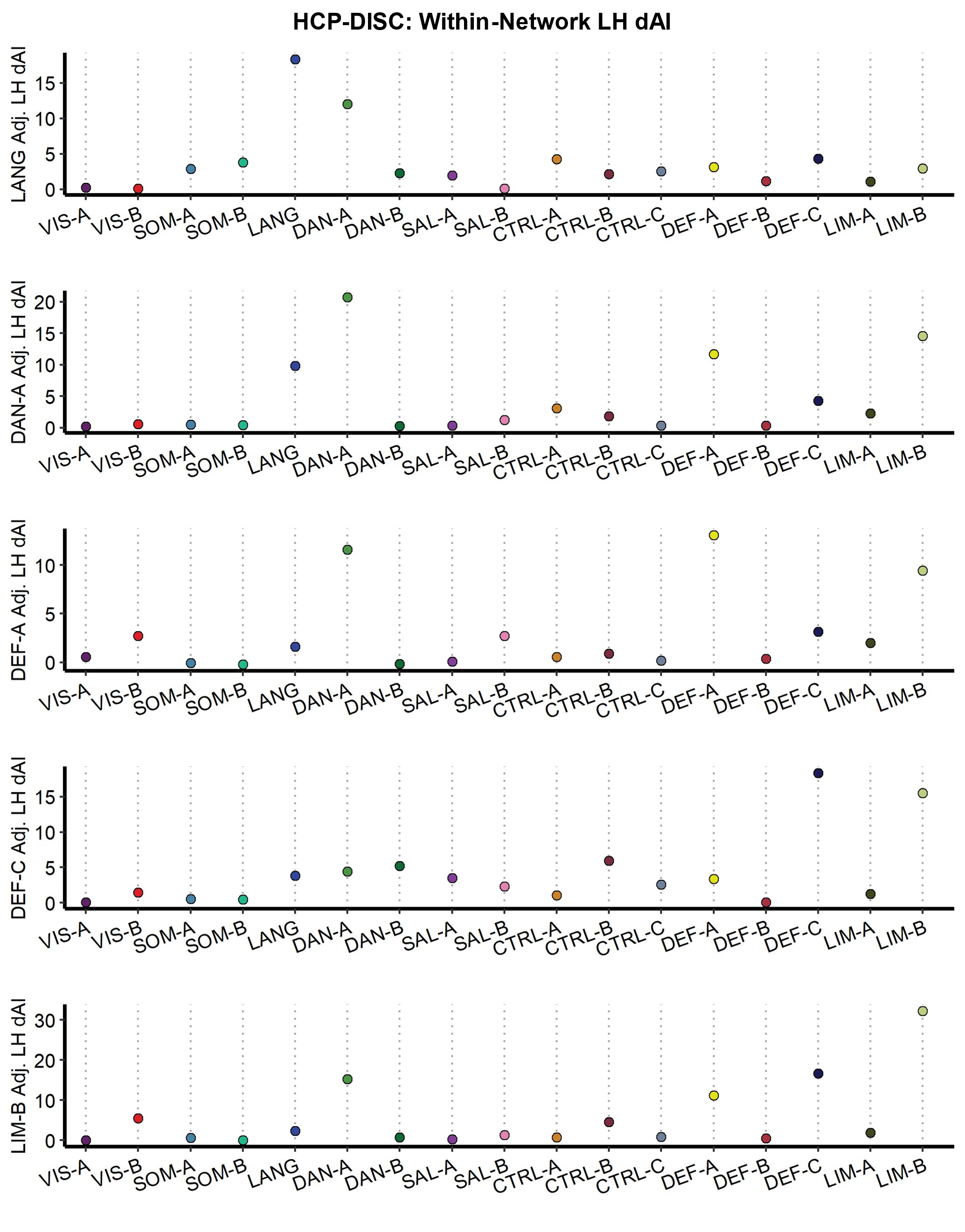
 **Figure S5.** Deconstructed autonomy index (dAI) for the top five left-lateralized networks in HCP-Discovery dataset. dAI values were averaged within each network (1-17) for each target network (language, dorsal attention-A, default-A, default-C, and limbic-B). Points represent mean dAI values.

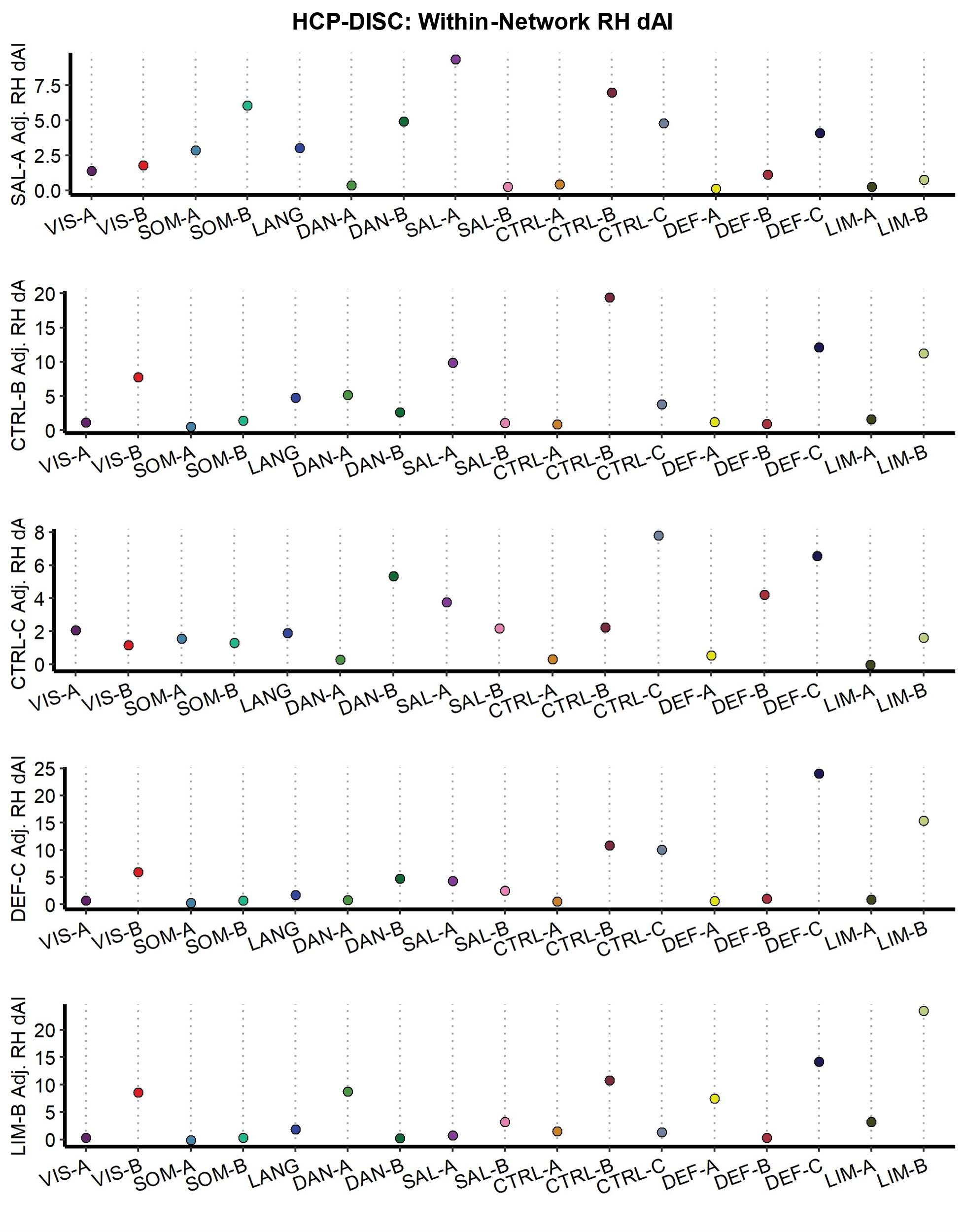
 **Figure S6.** Deconstructed autonomy index (dAI) for the top five right-lateralized networks in HCP-Discovery dataset. dAI values were averaged within each network (1-17) for each target network (salience/ventral attention-A, control-B, control-C, default-C, and limbic-B). Points represent mean dAI values.

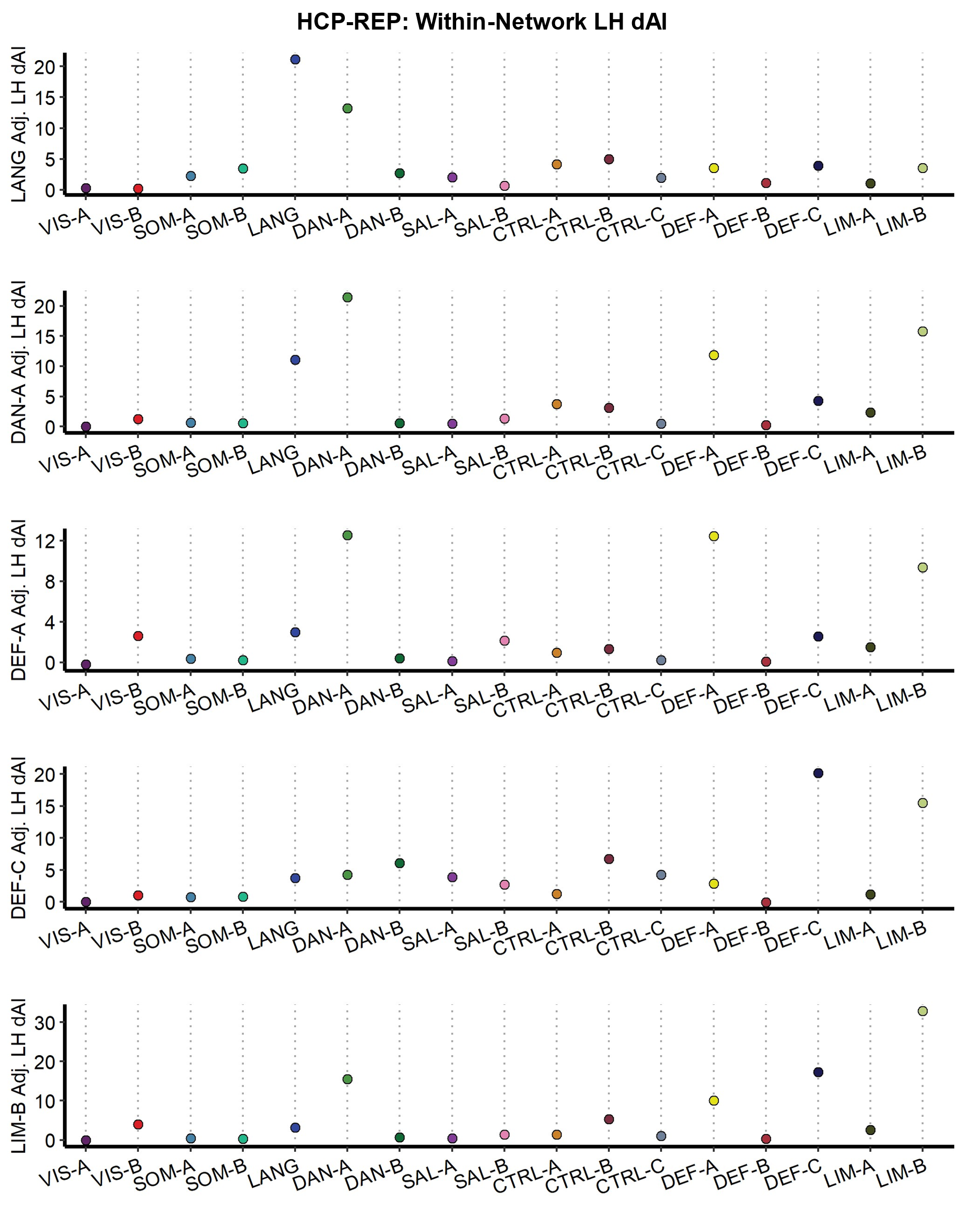
 **Figure S7.** Deconstructed autonomy index (dAI) for the top five left-lateralized networks in HCP-Replication dataset. dAI values were averaged within each network (1-17) for each target network (language, dorsal attention-A, default-A, default-C, and limbic-B). Points represent mean dAI values.

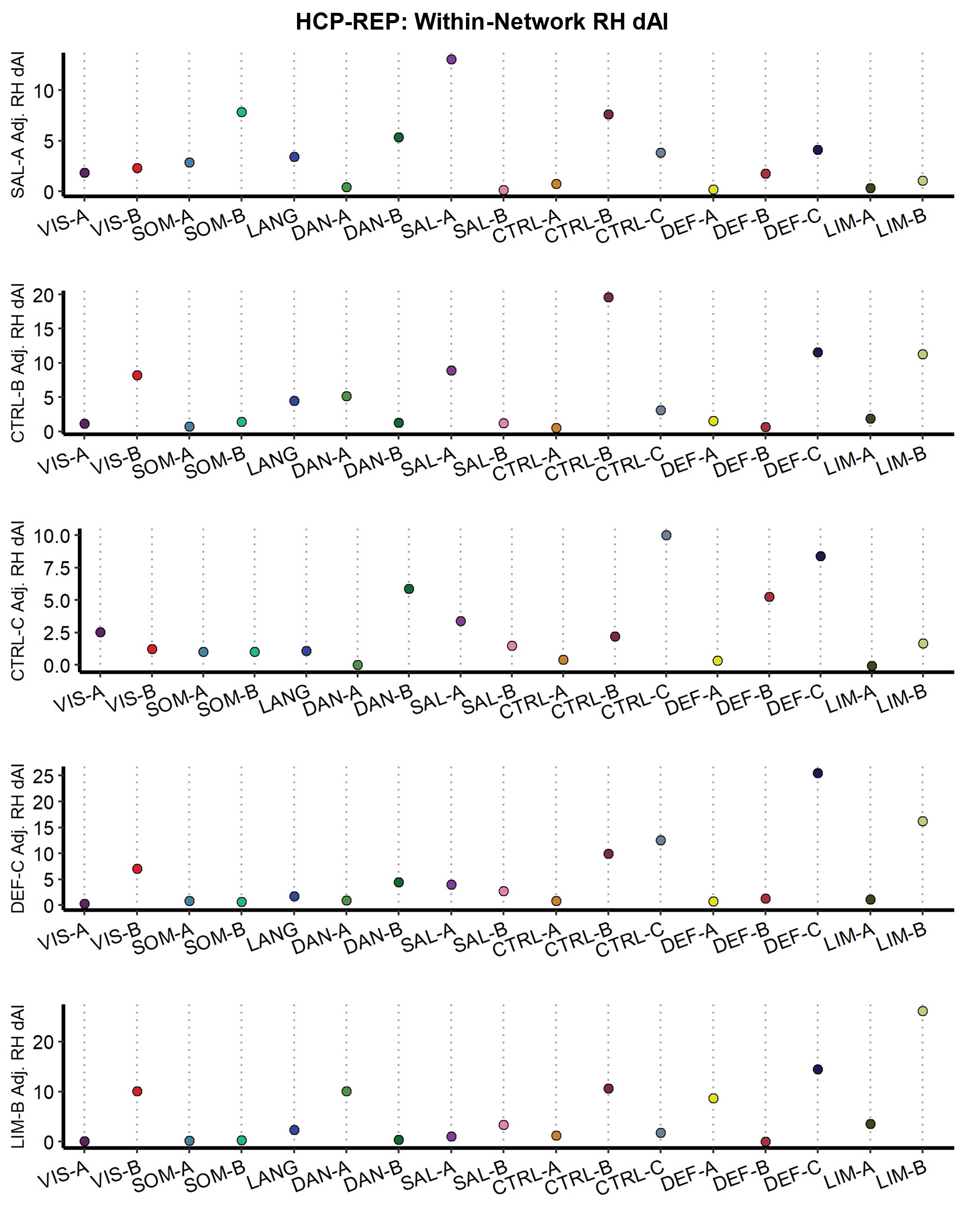

**Figure S8.** Deconstructed autonomy index (dAI) for the top five right-lateralized networks in HCP-Replication dataset. dAI values were averaged within each network (1-17) for each target network (salience/ventral attention-A, control-B, control-C, default-C, and limbic-B). Points represent mean dAI values.

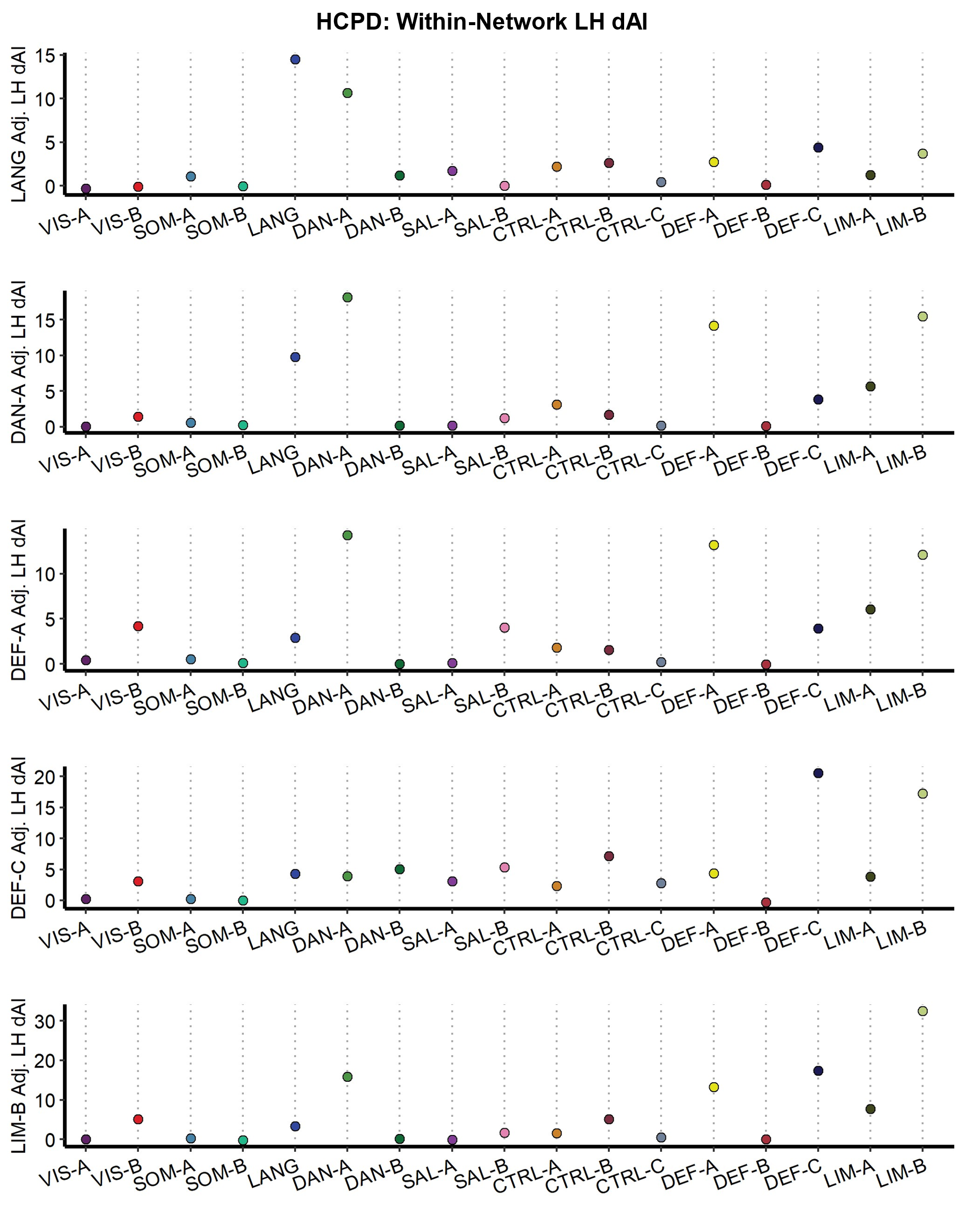
 **Figure S9.** Deconstructed autonomy index (dAI) for the top five left-lateralized networks in HCPD dataset. dAI values were averaged within each network (1-17) for each target network (language, dorsal attention-A, default-A, default-C, and limbic-B). Points represent mean dAI values.

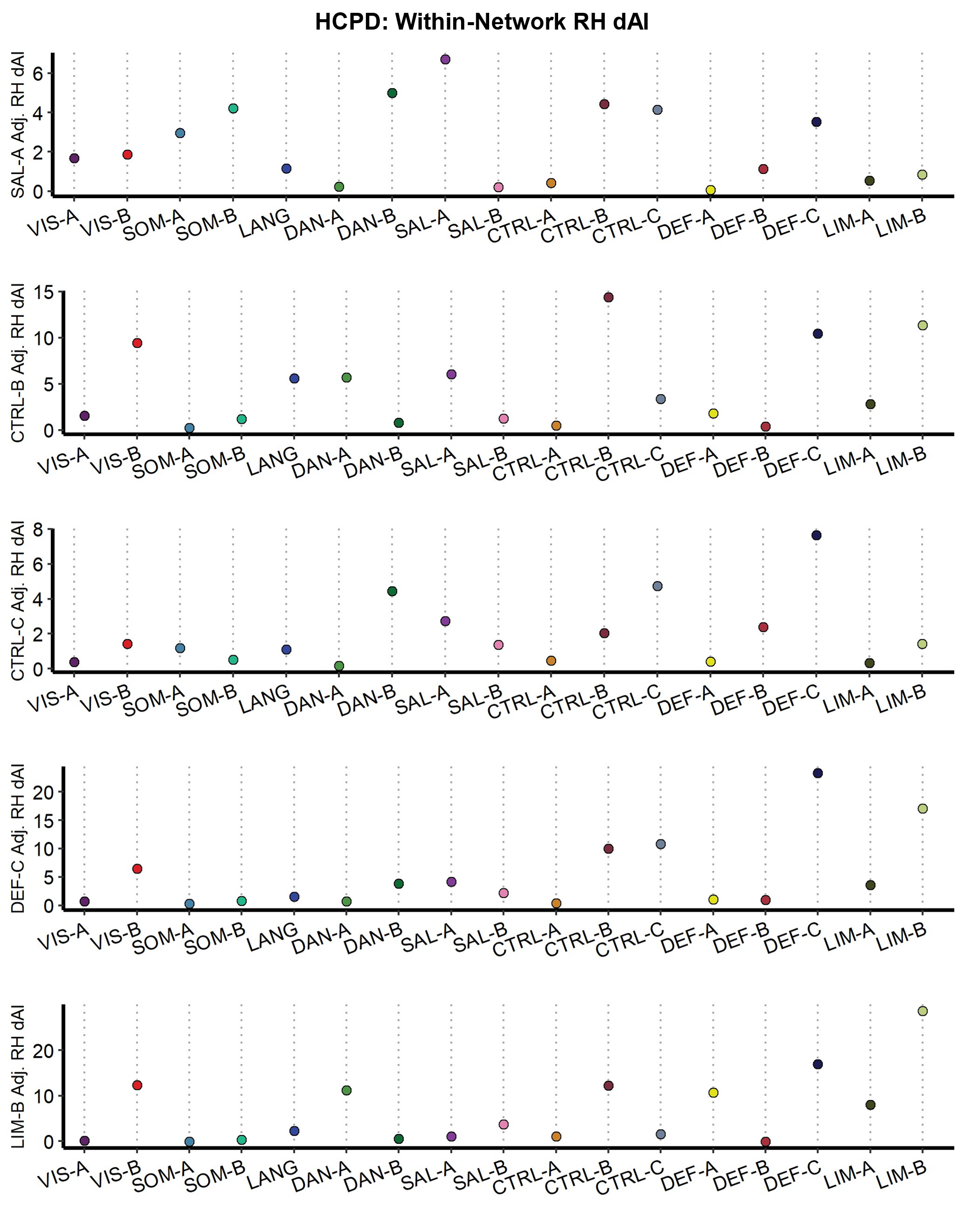

**Figure S10.** Deconstructed autonomy index (dAI) for the top five right-lateralized networks in HCPD dataset. dAI values were averaged within each network (1-17) for each target network (salience/ventral attention-A, control-B, control-C, default-C, and limbic-B). Points represent mean dAI values.

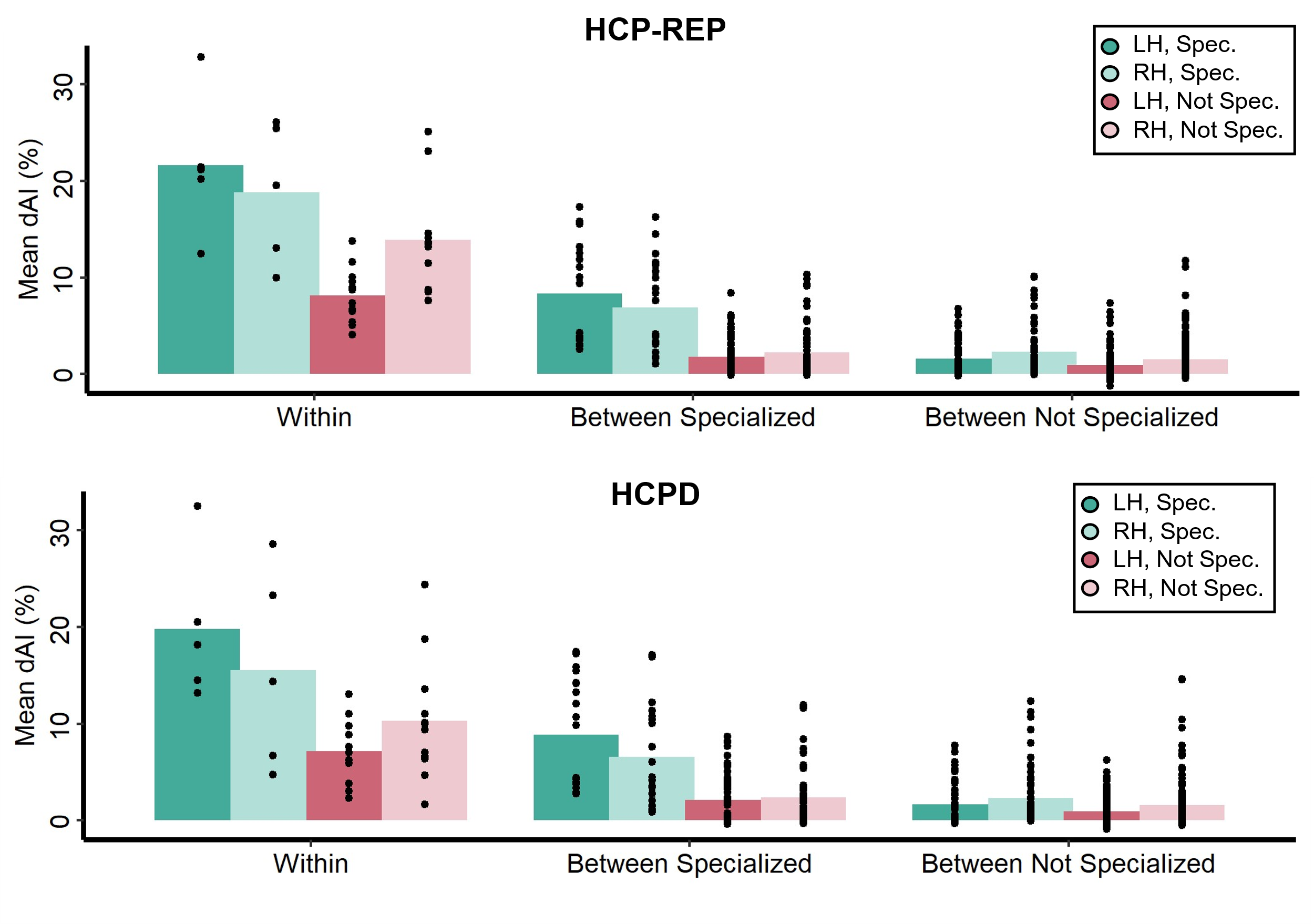
**Figure S11.** The deconstructed autonomy index in the HCP-Replication and HCPD datasets. For each row, the averaged dAI value for each category is given. dAI scores were grouped as being within-network (e.g., target network language and averaged network language), between specialized networks (e.g., target network language and averaged network dorsal attention-A), and between not specialized (e.g., target network language and averaged network visual-A). Specialized networks are the top five left- and right-lateralized networks (indicated with black boxes in Figure 4). Next, dAI scores were further binned depending on the target network as being specialized or not specialized (specialized networks included the top five left- and right-lateralized networks). Finally, dAI scores were organized by hemisphere, left or right. Each point represents a single target and averaged network combination mean dAI score. Across each dataset, within-network contributions appear to be greatest followed by between-specialized network contributions.

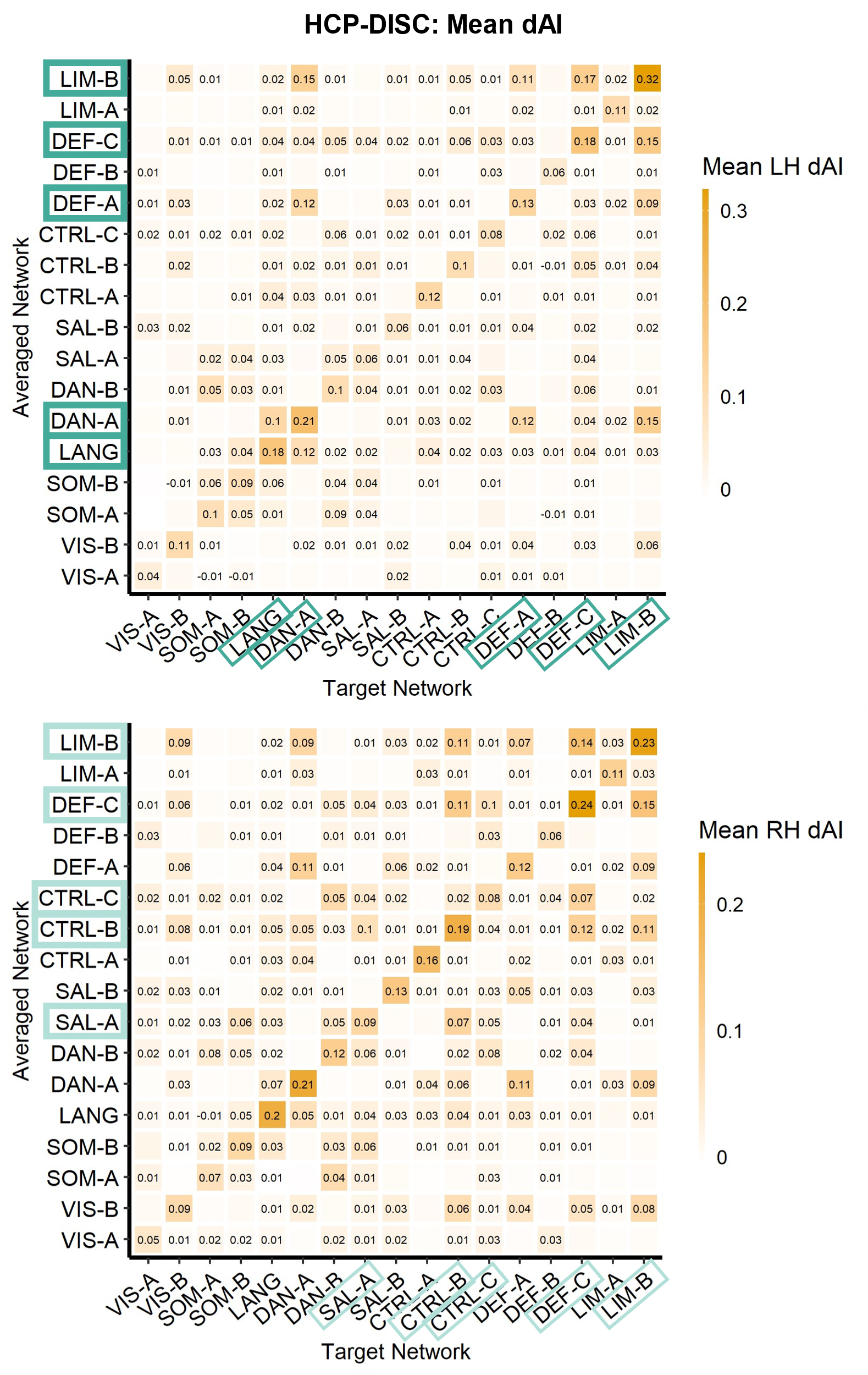
 **Figure S12.** Deconstructed autonomy index (dAI) for all 17 target networks in the HCP-Discovery dataset. dAI values were averaged within each network (1-17) for each target network (1-17). The diagonal represents within-network contributions, which appear to make the strongest contributions to network specialization. These are followed by contributions from other specialized networks.

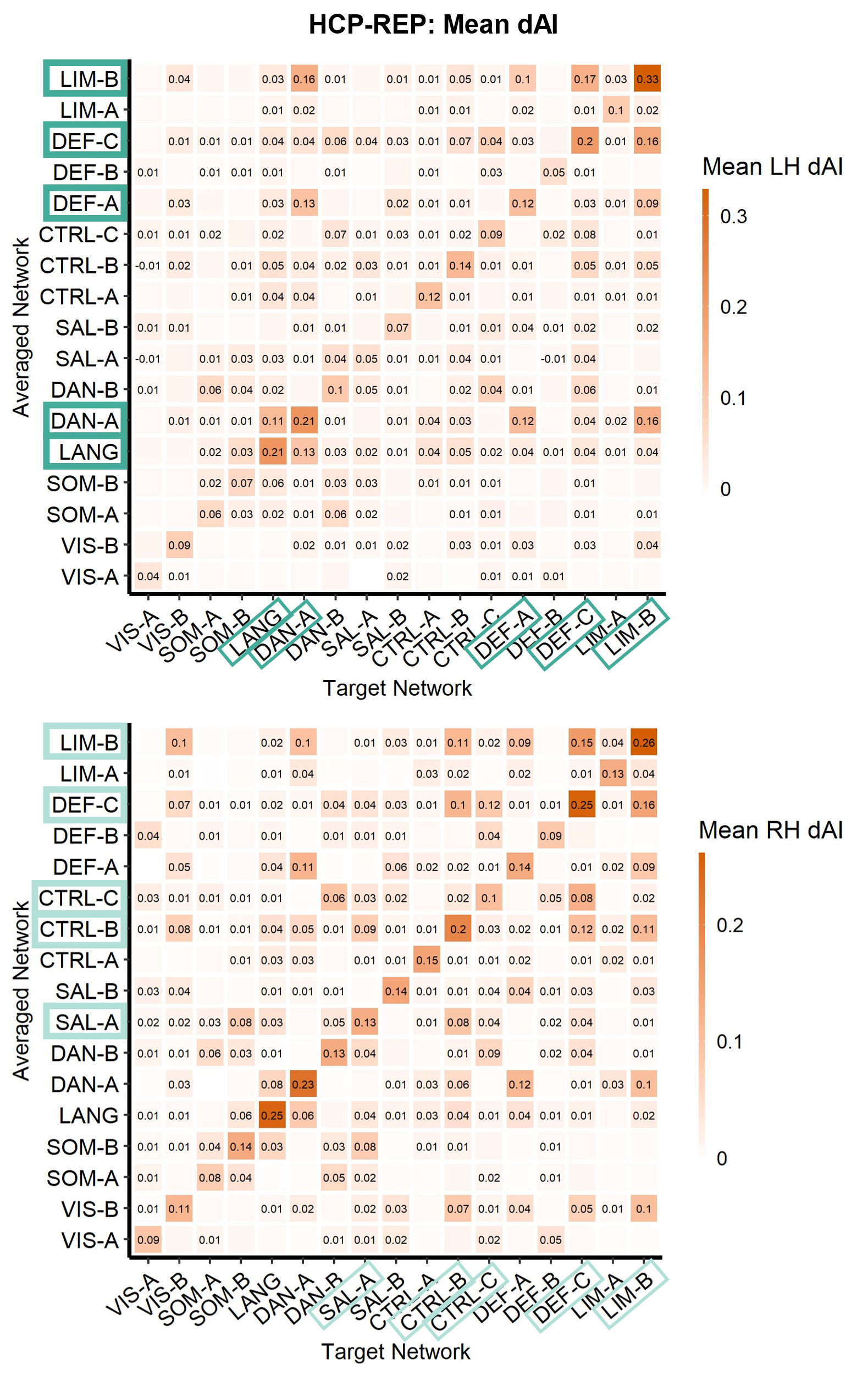
 **Figure S13.** Deconstructed autonomy index (dAI) for all 17 seed networks in the HCP-Replication dataset. dAI values were averaged within each network (1-17) for each target network (1-17). The diagonal represents within-network contributions, which appear to make the strongest contributions to network specialization. These are followed by contributions from other specialized networks.

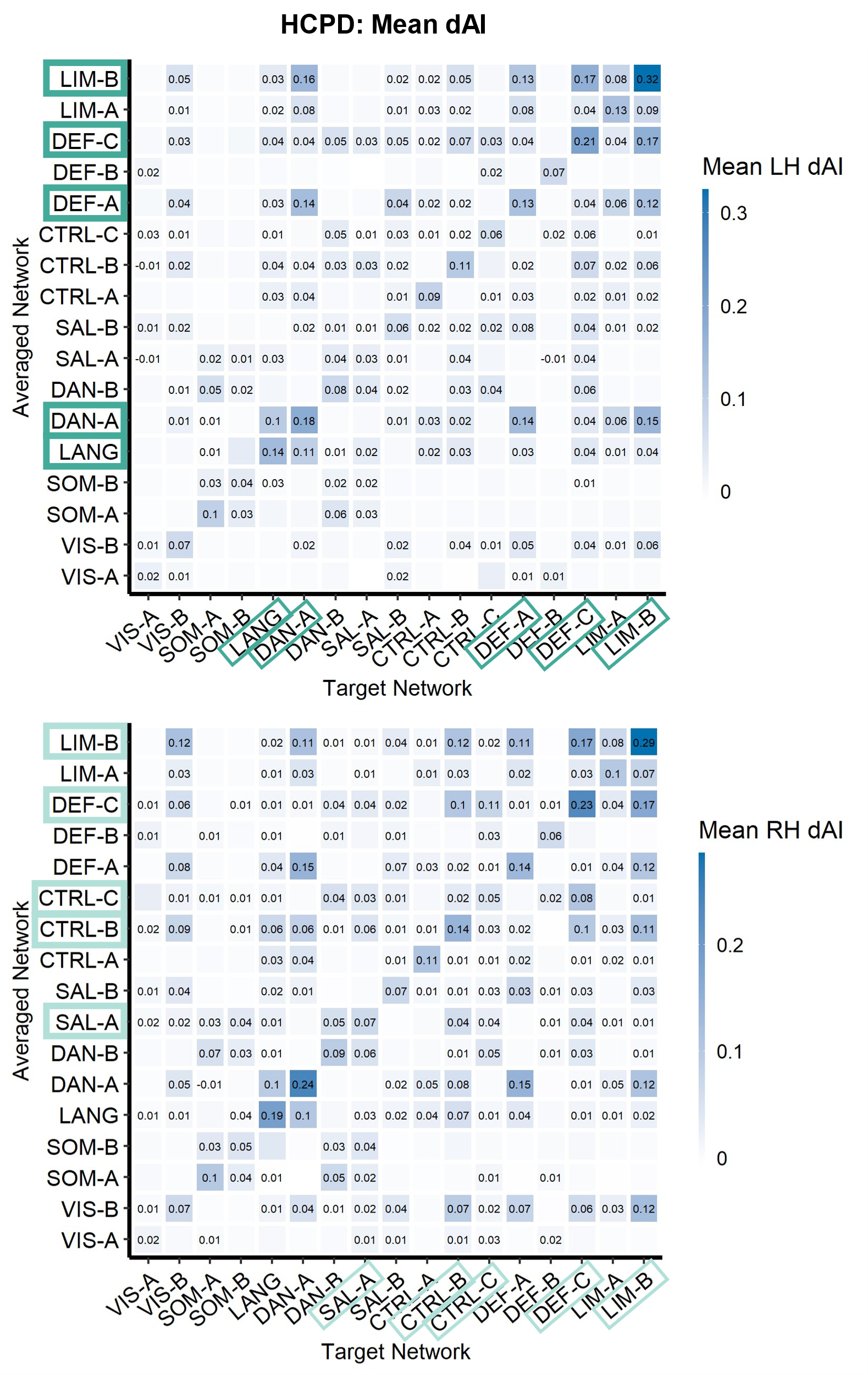
 **Figure S14.** Deconstructed autonomy index (dAI) for all 17 seed networks in the HCPD dataset. dAI values were averaged within each network (1-17) for each target network (1-17). The diagonal represents within-network contributions, which appear to make the strongest contributions to network specialization. These are followed by contributions from other specialized networks.

**Supplementary Table 1**

*Identifying Left-Specialized Networks Using Multiple Regressions in the HCP-Discovery Dataset (N = 276), HCP-Replication Dataset (N = 277), and the HCPD Dataset (N = 343)*

| Network Intercept | Dataset | β | *SE* | *t* | *p* |
| --- | --- | --- | --- | --- | --- |
| Visual-A |  |  |  |  |  |
|  | HCP-DISC | 0.19 | 0.18 | 1.07 | .29 |
|  | HCPD-REP | 0.01 | 0.02 | 0.29 | .14 |
|  | HCPD | 0.11 | 0.13 | 0.84 | .39 |
| **Visual-B** |  |  |  |  |  |
|  | HCP-DISC | 0.99 | 0.16 | 6.4 | < .001 |
|  | HCPD-REP | 0.75 | 0.15 | 5.06 | < .001 |
|  | HCPD | 0.97 | 0.18 | 5.3 | < .001 |
| **Somatomotor-A** |  |  |  |  |  |
|  | HCP-DISC | 1.66 | 0.21 | 7.52 | < .001 |
|  | HCPD-REP | 1.44 | 0.23 | 6.28 | < .001 |
|  | HCPD | 1.3 | 0.14 | 9.33 | < .001 |
| **Somatomotor-B** |  |  |  |  |  |
|  | HCP-DISC | 1.78 | 0.24 | 7.48 | < .001 |
|  | HCPD-REP | 1.79 | 0.19 | -0.68 | < .001 |
|  | HCPD | 0.63 | 0.12 | 5.3 | < .001 |
| **Language** |  |  |  |  |  |
|  | HCP-DISC | 3.53 | 0.32 | 11.12 | < .001 |
|  | HCPD-REP | 4.22 | 0.32 | 13.16 | < .001 |
|  | HCPD | 3.87 | 0.21 | 18.14 | < .001 |
| **Dorsal Attention-A** |  |  |  |  |  |
|  | HCP-DISC | 5.13 | 0.22 | 23.79 | < .001 |
|  | HCPD-REP | 5.79 | 0.24 | 24.09 | < .001 |
|  | HCPD | 5.89 | 0.22 | 27.09 | < .001 |
| **Dorsal Attention-B** |  |  |  |  |  |
|  | HCP-DISC | 2.57 | 0.22 | 11.44 | < .001 |
|  | HCPD-REP | 2.7 | 0.22 | 12.09 | < .001 |
|  | HCPD | 1.73 | 0.18 | 9.81 | < .001 |
| **Salience/VenAttn-A** |  |  |  |  |  |
|  | HCP-DISC | 1.55 | 0.23 | 6.76 | < .001 |
|  | HCPD-REP | 1.58 | 0.22 | 7.29 | < .001 |
|  | HCPD | 0.76 | 0.17 | 4.38 | < .001 |
| **Salience/VenAttn-B** |  |  |  |  |  |
|  | HCP-DISC | 1.07 | 0.13 | 7.89 | < .001 |
|  | HCPD-REP | 1.24 | 0.15 | 8.36 | < .001 |
|  | HCPD | 1.77 | 0.16 | 11.28 | < .001 |
| **Control-A** |  |  |  |  |  |
|  | HCP-DISC | 1.32 | 0.18 | 7.14 | < .001 |
|  | HCPD-REP | 1.59 | 0.16 | 9.79 | < .001 |
|  | HCPD | 1.57 | 0.09 | 16.75 | < .001 |
| **Control-B** |  |  |  |  |  |
|  | HCP-DISC | 1.56 | 0.24 | 6.46 | < .001 |
|  | HCPD-REP | 1.97 | 0.24 | 8.36 | < .001 |
|  | HCPD | 1.94 | 0.19 | 9.91 | < .001 |
| **Control-C** |  |  |  |  |  |
|  | HCP-DISC | 1.19 | 0.23 | 5.19 | < .001 |
|  | HCPD-REP | 1.25 | 0.21 | 6.07 | < .001 |
|  | HCPD | 0.58 | 0.18 | 3.28 | 0.001 |
| **Default-A** |  |  |  |  |  |
|  | HCP-DISC | 3.16 | 0.2 | 15.54 | < .001 |
|  | HCPD-REP | 3.59 | 0.22 | 16.67 | < .001 |
|  | HCPD | 4.42 | 0.21 | 20.78 | < .001 |
| Default-B |  |  |  |  |  |
|  | HCP-DISC | 0.4 | 0.16 | 2.57 | 0.01 |
|  | HCPD-REP | 0.39 | 0.17 | 2.24 | 0.03 |
|  | HCPD | 0.14 | 0.13 | 1.07 | 0.28 |
| **Default-C** |  |  |  |  |  |
|  | HCP-DISC | 4.06 | 0.21 | 19.47 | < .001 |
|  | HCPD-REP | 3.85 | 0.2 | 18.91 | < .001 |
|  | HCPD | 4.33 | 0.2 | 21.17 | < .001 |
| **Limbic-A** |  |  |  |  |  |
|  | HCP-DISC | 0.92 | 0.09 | 10.09 | < .001 |
|  | HCPD-REP | 1.01 | 0.11 | 9.26 | < .001 |
|  | HCPD | 2.12 | 0.13 | 16.16 | < .001 |
| **Limbic-B** |  |  |  |  |  |
|  | HCP-DISC | 4.11 | 0.21 | 19.27 | < .001 |
|  | HCPD-REP | 3.91 | 0.22 | 18.09 | < .001 |
|  | HCPD | 4.7 | 0.21 | 22.31 | < .001 |

*Note:* Coefficients and *p*-values for the intercept are shown. Networks with reliably significant (Bonferroni-corrected alpha level of .001) intercepts are bolded.

**Supplementary Table 2**

*Identifying Right-Specialized Networks Using Multiple Regressions in the HCP-Discovery Dataset (N = 276), HCP-Replication Dataset (N = 277), and the HCPD Dataset (N = 343)*

| Network Intercept | Dataset | β | *SE* | *t* | *p* |
| --- | --- | --- | --- | --- | --- |
| **Visual-A** |  |  |  |  |  |
|  | HCP-DISC | 1.39 | 0.18 | 7.69 | < .001 |
|  | HCPD-REP | 1.52 | 0.19 | 7.76 | < .001 |
|  | HCPD | 0.93 | 0.14 | 6.72 | < .001 |
| **Visual-B** |  |  |  |  |  |
|  | HCP-DISC | 2.4 | 0.2 | 11.97 | < .001 |
|  | HCPD-REP | 2.79 | 0.21 | 13.48 | < .001 |
|  | HCPD | 3.27 | 0.21 | 15.89 | < .001 |
| **Somatomotor-A** |  |  |  |  |  |
|  | HCP-DISC | 1.84 | 0.2 | 9 | < .001 |
|  | HCPD-REP | 1.86 | 0.2 | 9.29 | < .001 |
|  | HCPD | 1.78 | 0.13 | 13.46 | < .001 |
| **Somatomotor-B** |  |  |  |  |  |
|  | HCP-DISC | 2.02 | 0.18 | 11.07 | < .001 |
|  | HCPD-REP | 2.33 | 0.17 | 14.01 | < .001 |
|  | HCPD | 1.46 | 0.13 | 11.64 | < .001 |
| **Language** |  |  |  |  |  |
|  | HCP-DISC | 2.41 | 0.18 | 13.45 | < .001 |
|  | HCPD-REP | 2.75 | 0.21 | 13.03 | < .001 |
|  | HCPD | 1.61 | 0.18 | 8.91 | < .001 |
| **Dorsal Attention-A** |  |  |  |  |  |
|  | HCP-DISC | 2.15 | 0.19 | 11.32 | < .001 |
|  | HCPD-REP | 1.84 | 0.19 | 9.93 | < .001 |
|  | HCPD | 2.12 | 0.19 | 10.72 | < .001 |
| **Dorsal Attention-B** |  |  |  |  |  |
|  | HCP-DISC | 2.76 | 0.22 | 12.56 | < .001 |
|  | HCPD-REP | 2.67 | 0.24 | 11.15 | < .001 |
|  | HCPD | 2.54 | 0.19 | 13.21 | < .001 |
| **Salience/VenAttn-A** |  |  |  |  |  |
|  | HCP-DISC | 2.83 | 0.24 | 11.82 | < .001 |
|  | HCPD-REP | 3.14 | 0.26 | 11.95 | < .001 |
|  | HCPD | 2.74 | 0.21 | 13.05 | < .001 |
| **Salience/VenAttn-B** |  |  |  |  |  |
|  | HCP-DISC | 1.73 | 0.14 | 12.61 | < .001 |
|  | HCPD-REP | 1.75 | 0.14 | 12.45 | < .001 |
|  | HCPD | 1.49 | 0.14 | 10.63 | < .001 |
| **Control-A** |  |  |  |  |  |
|  | HCP-DISC | 1.22 | 0.11 | 11.56 | < .001 |
|  | HCPD-REP | 1.24 | 0.1 | 11.9 | < .001 |
|  | HCPD | 0.84 | 0.06 | 13.14 | < .001 |
| **Control-B** |  |  |  |  |  |
|  | HCP-DISC | 4.09 | 0.26 | 15.68 | < .001 |
|  | HCPD-REP | 4.09 | 0.26 | 15.79 | < .001 |
|  | HCPD | 4.22 | 0.24 | 17.89 | < .001 |
| **Control-C** |  |  |  |  |  |
|  | HCP-DISC | 2.91 | 0.22 | 13.03 | < .001 |
|  | HCPD-REP | 3.14 | 0.23 | 13.67 | < .001 |
|  | HCPD | 2.84 | 0.19 | 15.15 | 0.001 |
| **Default-A** |  |  |  |  |  |
|  | HCP-DISC | 1.98 | 0.21 | 9.64 | < .001 |
|  | HCPD-REP | 1.83 | 0.18 | 9.95 | < .001 |
|  | HCPD | 1.97 | 0.19 | 9.98 | < .001 |
| **Default-B** |  |  |  |  |  |
|  | HCP-DISC | 1.46 | 0.17 | 8.51 | < .001 |
|  | HCPD-REP | 1.91 | 0.19 | 9.87 | < .001 |
|  | HCPD | 1.19 | 0.14 | 8.38 | < .001 |
| **Default-C** |  |  |  |  |  |
|  | HCP-DISC | 3.62 | 0.22 | 16.21 | < .001 |
|  | HCPD-REP | 4.12 | 0.23 | 17.92 | < .001 |
|  | HCPD | 4.33 | 0.2 | 21.44 | < .001 |
| **Limbic-A** |  |  |  |  |  |
|  | HCP-DISC | 1.14 | 0.15 | 7.85 | < .001 |
|  | HCPD-REP | 1.24 | 0.14 | 9.16 | < .001 |
|  | HCPD | 2.01 | 0.11 | 17.59 | < .001 |
| **Limbic-B** |  |  |  |  |  |
|  | HCP-DISC | 5.06 | 0.26 | 19.69 | < .001 |
|  | HCPD-REP | 5.7 | 0.23 | 24.29 | < .001 |
|  | HCPD | 6.07 | 0.22 | 27.85 | < .001 |

*Note:* Coefficients and *p*-values for the intercept are shown. Networks with reliably significant (Bonferroni-corrected alpha level of .001) intercepts are bolded.
